# mDrop-seq: Massively parallel single-cell RNA-seq of *Saccharomyces cerevisiae* and *Candida albicans*

**DOI:** 10.1101/2021.07.06.451338

**Authors:** Ryan Dohn, Bingqing Xie, Rebecca Back, Alan Selewa, Heather Eckart, Reeta Prusty Rao, A. Basu

**Affiliations:** Section of Genetic Medicine, Department of Medicine, University of Chicago; Committee on Genetics, Genomics and Systems Biology, University of Chicago; Biophysical Sciences Graduate Program, University of Chicago; Department of Biology and Biotechnology, Worcester Polytechnic Institute

## Abstract

Advances in high-throughput single-cell mRNA sequencing (scRNA-seq) have been limited till date by technical challenges like tough cell walls and low RNA quantity that prevented transcriptomic profiling of microbial species at throughput. We present microbial Drop-seq or mDrop-seq, a high-throughput scRNA-seq technique that is used on two yeast species, *Saccharomyces cerevisiae*, a popular model organism and *Candida albicans*, a common opportunistic pathogen. We benchmarked mDrop-seq for sensitivity and specificity and used it to profile 35,109 *S. cerevisiae* cells to detect variation in mRNA levels between them. As a proof of concept, we quantified expression differences in heat-shocked *S. cerevisiae* using mDrop-seq. We detected differential activation of stress response genes within a seemingly homogenous population of *S. cerevisiae* under heat-shock. We also applied mDrop-seq to *C. albicans* cells, a polymorphic and clinically relevant yeast species with thicker cell wall compared to *S. cerevisiae*. Single cell transcriptomes in 39,705 *C. albicans* cells was characterized using mDrop-seq under different conditions, including exposure to fluconazole, a common anti-fungal drug. We noted differential regulation in stress response and drug target pathways between *C. albicans* cells, changes in cell cycle patterns and marked increases in histone activity. These experiments are among the first high throughput single cell RNA-seq on different yeast species and demonstrate mDrop-seq as an affordable, easily implementable, and scalable technique that can quantify the variability in gene expression in different yeast species. We hope that mDrop-seq will lead to better understanding of genetic variation in yeasts in response to stimuli and find immediate applications in investigating drug resistance and infection outcome.

## Introduction

The rise of high-throughput single-cell mRNA sequencing (scRNA-seq) has led to greater understanding of the functional and phenotypic heterogeneity present in our body on a cellular level. Primarily developed for mammalian cells [1], scRNA-seq uses the transcriptome of a single cell to analyze of cell type [1], cell state [2], and cell response.[3] While variation between different cell types (in a multicellular organism) or cells of different species may be expected, scRNA-seq techniques have shown that there is significant cell-to-cell heterogeneity even between otherwise identical cells.[4] High-throughput techniques can examine thousands of cells at once, adding statistical power to determine variability between cells.[1], [5]

Technological challenges, such as the tough cell walls, small size, and concomitantly smaller amounts of transcripts per cell [6] however, have prevented similar applications in unicellular microbial organisms.[7] The strength and rigidity of the microbial cell walls composed of diverse components, e.g., peptidoglycans in bacteria, and chitin and β-glucan layers in yeasts [8]–[10] make them resistant to most lysis agents used for scRNA-seq. Microbes also have significantly less mRNA compared to animal cells, with estimates ranging up to two orders of magnitude less for bacterial cells.[11], [12]

Despite the challenges, achieving high-throughput microbial scRNA-seq is essential to understanding the heterogeneity and complex interactions between cells in a population. A few recent studies profiled *S. cerevisiae* at single cell resolution [13], [14], [15]. However, clinically relevant yeast species, such as *Candida albicans*, have yet to be characterized with single-cell resolution in a high-throughput manner. Single-cell RNA-seq on fungal pathogens such as *Candida albicans* and *Candida auris* can help understand the commensal-to-pathogenic switch that lead to opportunistic infections.[16], [17] As a common hospital acquired infection, *C. albicans* can cause both superficial infections in humans and severe systemic infections in immunocompromised individuals.[18] Understanding the heterogeneity in how individual yeast cells respond to changes in the hosts’ immune system or their shifting microbiome can help treat and prevent such infections. The emergence of drug resistant microbes is an urgent human health crisis.[19] Molecular mechanisms that confer protection to a select few cells when the parent population is decimated by anti-microbial agents can help understand the rise of drug-resistant strains.[20]

Here, we present microbial Drop-seq, or mDrop-seq as a method of high-throughput scRNA-seq of different yeast species. With modifications to the Drop-seq platform [1], we were able to accomplish microfluidic compartmentalization of single cells, in-droplet lysis and cellular barcoding of two species of yeast, *viz*., *S. cerevisiae* and *C. albicans* for scRNA-seq at scale. We quantified the transcriptional heterogeneity within clonal populations of yeast cells and profiled their response at single-cell resolution to environmental stress, such as heat shock, and exposure to fluconazole, a common anti-fungal agent. Under heat shock at 42 °C, we observe differential expression of key stress response genes, including *HSP12, HSP26*, and *HSP42* in *S. cerevisiae* cells. When exposed to fluconazole, *C. albicans* cells show differential upregulation of key drug target pathways such as ergosterol biosynthesis and ribosome activity, and an overall increase in histone activity. Importantly, both *S. cerevisiae* and *C. albicans* show disruption in their cell cycle patterns under heat shock and fluconazole exposure, respectively. Taken together, we posit that the cell cycle state of the yeast cell in a population of continuously cycling cells is related to the variability in the cell’s response to stress. mDrop-seq’s ability to simultaneously profile a mix two phenotypically different species demonstrate that different fungal species are amenable to single-cell transcriptomic profiling using mDrop-seq. scRNA-seq of a mix of *S. cerevisiae* and *C. albicans* also demonstrates that mDrop-seq is capable of simultaneous single-cell profiling of a population of different yeast species.

## Results

### Optimizing single yeast cell lysis in droplets

To establish a high-throughput scRNA-seq method for yeast cells such as *S. cerevisiae* and *C. albicans*, we adapted the emulsion droplet and bead-based DNA barcoding scheme previously used in Drop-seq [1] and DroNc-seq.[21] Initial experiments were performed using *S. cerevisiae*, a popular model organism that is amenable to technology development due to the species’ relatively thinner cell wall (∼120 nm, compared to ∼150 nm thick cell walls in *C. albicans* [22]), and easier lysis. *S. cerevisiae* also has a well-annotated genome that is useful to establish data analysis tools. Laboratory strains BY4741 and SC5314 were used for *S. cerevisiae* and *C. albicans*, respectively.

The scRNA-seq workflow in yeasts in microfluidic drops is summarized in Figure 1A. To determine the duration of droplet incubation for optimal zymolyase activity needed for each species, a single collection of mDrop-seq droplets on *S. cerevisiae* was split into three different pools and each pool was incubated in a 37 °C water bath for 10, 15, and 20 minutes, guided by bulk lysis experiments. The droplets were inspected by optical microscopy following incubation, to ensure droplet stability and cell lysis.

**Figure 1:**
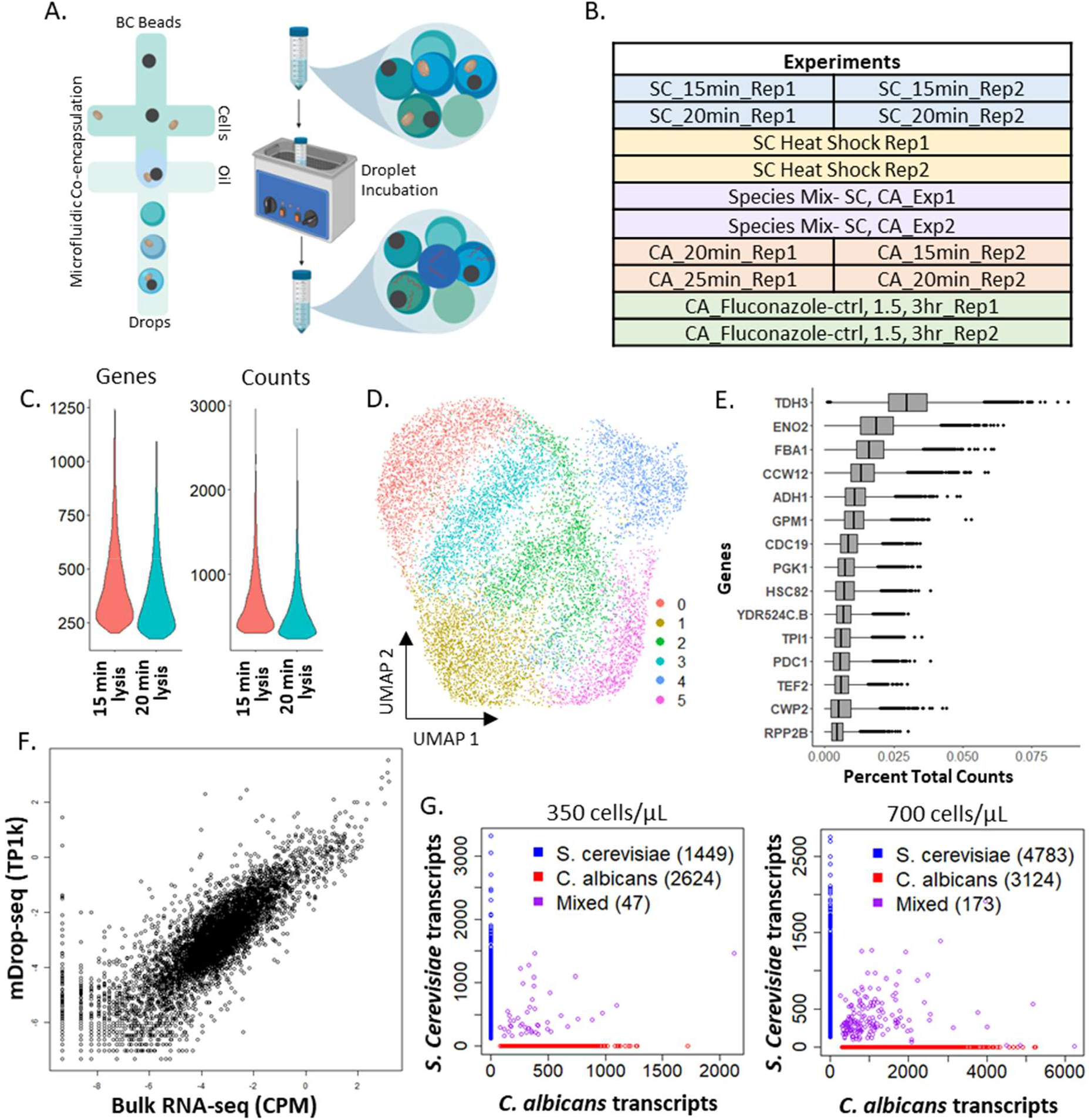
mDrop-seq of *Saccharomyces cerevisiae* cells. A) mDrop-seq experimental schematic. B) List of mDrop-seq experiments on different yeast samples. C) Violin plots showing the number of genes and UMI detected for each cell at two cell lysis times: 15 and 20 minutes. D) UMAP visualizing the results of clustering analysis on 12,012 *S. cerevisiae* cells. E) Boxplot displaying the top 15 genes expressed by percentage of total counts across 15 and 20 min lysis times. F) Correlation plot of average gene expression between mDrop-seq (SC_15min_rep1) and bulk RNA-seq. G) Species-mixing plots where each dot depicts a unique cellular barcode that align to *S. cerevisiae* (blue), *C. albicans* (red), or both genomes (purple). Two Poisson loading concentrations of 300 cells/µL (left; λ = 0.08) and 700 cells/µL (right; λ = 0.15) are tested.

The 10-min lysis incubation yielded lower quantity of cDNA, indicating incomplete lysis and were excluded from further analysis. The 15 and 20 min incubations generated two libraries, indicated as SC_15min_Rep1 and SC_20min_Rep1. There were ∼5000-10,000 cells processed per sample. Figure 1C shows the number of features (left), as a proxy for genes, and Unique Molecular Identifiers or UMI (right), as a proxy for number of mRNA molecules captured per cell barcode. The UMI identifies individual mRNA molecules detected, allowing correction for PCR replicates to prevent PCR bias. Supplementary Table S1 summarizes the total and average number of reads from each sample, number of cells, mean number of genes and UMI obtained from a single cell, and the total number of unique genes obtained per experiment. We filtered out cell barcodes with less than 200 or more than 2000 genes detected, as they likely represent poor quality cells or multiple cells loaded in a single droplet, respectively. Across all experiments, we saw only 30-35% more UMI compared to the number of genes detected [23]. This experiment was performed twice.

The two mDrop-seq datasets from the lysis trial, SC_15min_Rep1, SC_20min_Rep1 overlap in principal component (PC) space (Figures S1A, B, C), with a Pearson correlation > 0.99 between the datasets for UMI counts. This allowed us to combine the SC_15min_Rep1, SC_20min_Rep1 datasets into a single dataset of 12,012 cells which we used to systematically explore differential expression (DE) in *S. cerevisiae* genes. Figure 1D shows the combined dataset in Uniform Manifold Approximation and Projection (UMAP) space.[24] The genes with highest expression in this dataset were involved in glycolysis (*TDH3, ENO2, FBA1*), stress response (*HSC82*), or ribosome biogenesis (*RPP2B*) (Figure 1E). As these cells were grown in 2% glucose medium and lysed in log phase, the presence of glycolysis genes is expected. We also saw expression of general stress response genes common to heat shock, DNA replication stress and oxidative stress that we attribute to the yeast cells responding to stresses (e.g., enzymatic lysis, heat) during cell lysis.

To compare mDrop-seq with bulk RNA-seq, we also performed population-level RNA-seq on *S. cerevisiae*. A pseudo-bulk [1] comparison of log normalized transcript counts in mDrop-seq showed a Pearson’s correlation of 0.85 with our bulk RNA-seq data, and overall good correlation (∼0.8, on average), with public RNA-seq datasets.[25] The correlation in transcript counts between our bulk RNA-seq and public datasets was 0.85, comparable to mDrop-seq.

### Single cell specificity of mDrop-seq confirmed by species mixing experiments

To establish that we have single-cell specificity in our experiments, species-mixing experiments [1] were performed using a mix of *S. cerevisiae* and *C. albicans* cells. Species-mixing experiments allow checking for ‘doublets’, or cell barcodes that capture two cells, assuming that a fraction of such drops will capture both *S. cerevisiae* and *C. albicans* cells and the corresponding barcode beads will yield cell barcodes that align to both *S. cerevisiae* and *C. albicans* genomes. The probability of finding one or more cells in a single drop may be estimated assuming a uniform concentration of cells in the loading medium following a Poisson distribution. The Poisson parameter, λ governs the cell distribution in drops and may be used to calculate the fraction of cell doublets, etc. in the experiment. Two different cell-loading conditions were tested at 350,000 and 700,000 cells/mL, representing Poisson loading parameter, λ = 0.077 and 0.15, respectively. Across 4,120 cells detected at λ = 0.077 or 7.7% Poisson loading, majority of the barcodes map to one genome only, with only 47 barcodes mapping to both *S. cerevisiae* and *C. albicans* genomes (Figure 1G, left). Assuming that cross-species doublets make up half of all doublets (a similar number of barcodes would contain two *S. cerevisiae* cells or two *C. albicans* cells), we estimate the doublet rate to be ∼2.3% of all barcodes detected. Figure 1G, right shows a similar species-mixing experiment at λ = 0.15 or 15% Poisson loading; across 8,080 cells detected, only 173 cells map to both genomes, giving a final doublet rate of ∼4.2%. A Poisson loading of 15% was used for all *S. cerevisiae* and *C. albicans* experiments. We note that scRNA-seq of *S. cerevisiae* and *C. albicans* in these species mixing experiments also demonstrate that mDrop-seq is capable of simultaneous single-cell profiling of a mix of different yeast species. This feature may be useful in characterizing fungal microbiome composed of multiple yeast species without the need to sort cells by species.

### Heterogeneity in heat shock response of *Saccharomyces cerevisiae* profiled using mDrop-seq

To test the efficacy of mDrop-seq in detecting transcriptional changes in yeast cells to external stimulus, *S. cerevisiae* cells were subjected to heat shock prior to running mDrop-seq. The heat shock is a widely conserved response in cells that involves expression of protein chaperones.[26] The cells underwent a 20-minute heat shock at 42 °C and then were chilled on ice before running mDrop-seq. The *S. cerevisiae* dataset shown previously in Figure 1D was used as the control dataset to the heat shock experiments.

Two replicate experiments on heat shock stimulation were performed. We see significant similarity in expression data with Pearson correlation of 0.96. The second replicate saw an increase in genes and UMI detected (587 and 1053, respectively), seen in Figure 2A. In PC space, the replicates cluster closely but separately from one another (Figure S4C). The mDrop-seq replicate experiments were performed on different days to characterize the heat shock response of *S. cerevisiae*, using independent cell cultures grown from the same stock.

**Figure 2:**
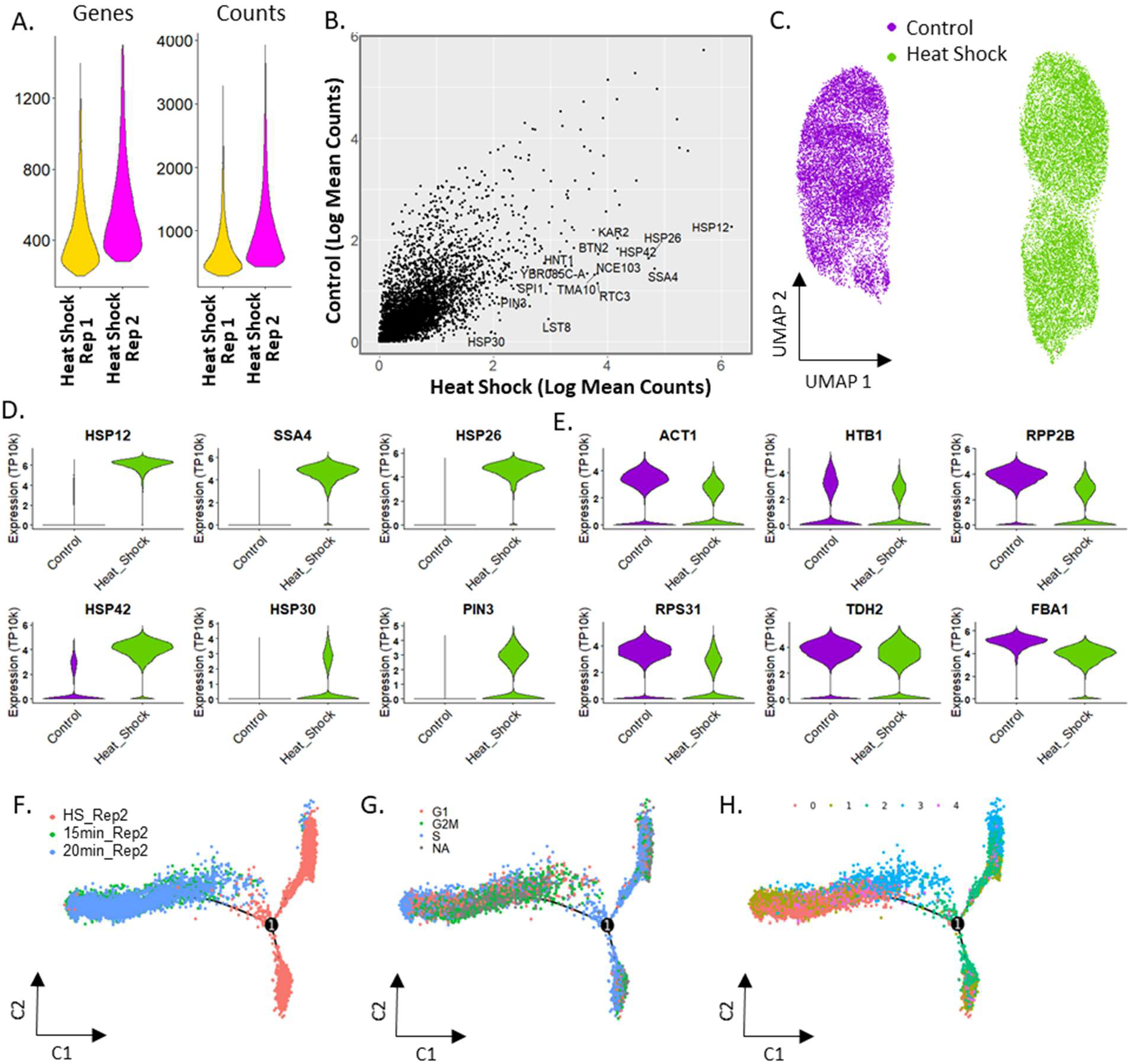
Heat shock treatment of 26,019 *Saccharomyces cerevisiae* cells profiled using mDrop-seq. A) Violin plot displaying the number of genes and UMI for each heat-shock replicate. B) Correlation between average gene expression values for the control and heat-shocked *S. cerevisiae* datasets. C) UMAP displaying the clustering patterns of the control and heat-shock datasets, including biological replicates. D) Violin plots displaying the expression differences in control and heat-shocked datasets for several heat-shock related genes. E) Violin plots displaying expression of several “house-keeping” genes (actin, histones, ribosomal, and glycolysis) in heat shock and control data. F-H) Pseudo-time trajectory of gene expression inferred from the combined control and heat shocked *S. cerevisiae* dataset. Colors indicate F) experimental time points, G) Cell cycle stages (cells that could not be assigned to a cell cycle stage are marked NA), and H) cell-type clusters shown in S3A.

Compared to the control, we see the upregulation of several stress response genes (the mean expression of the control and heat-shock data show Pearson correlation of 0.74; see Figure 2B). UMAP of the control and heat-shock datasets shows distinct clusters of heat-shocked and unstimulated cells (Figure 2C). While the heat-shock replicates cluster separately, they appear closer together on the UMAP, compared to the controls.

Next, we ordered the genes in the control and heat-shock datasets by descending log fold change (logFC). For many of these genes, the p-values are small (Wilcoxon Rank Sum test, adj-p < 1e-10, Bonferroni correction) with many adj-p < 2.225e-308 (lowest value reported on Seurat). Among the genes induced in cells under heat-shock compared to control, we see several genes involved in heat-shock related stress response, such as *HSP12* (logFC = 1.78) as well as other heat-shock protein (HSP) family genes associated with other types of stress response (Figure 2D). Protein transport genes, such as *KAR2* (logFC = 2.49), are also highly expressed under heat-shock (Figure S4A). We also see significant differential expression in genes marked for cell stress, such as *RTC3, NCE103* (logFC = 2.68, 2.51, respectively) involved in DNA replication stress (Figure S4B). House-keeping genes, such as actin (*ACT1*), histone genes (*HTB1*), ribosomal genes, and glycolysis genes (Figure 2E) are present in both datasets, with slightly higher expression (logFC = 0.6, 1.1, and 1.5 for *HTB2, ACT1*, and *RPP2B*, respectively) in the control dataset. These results suggest that mDrop-seq has the power to detect cellular responses to stimuli on a single cell level.

### Activation of stress response in *S. cerevisiae* under heat shock

To investigate the sequence of activation in stress response genes in SC under heat shock, we applied trajectory analysis [27] on a subset of control and heat-shock SC data. Three datasets, SC_15min_Rep2, SC_20min_Rep2 and SC_HeatShock_Rep2 were integrated (Figure S3A) and cell-cycle module scores were assigned to each cell, using Seurat. We then used Monocle v2 [27] to infer the expression changes during heat shock in pseudo-time (Figure 2F-H). Figure 2F shows the trajectory where the control samples, SC_15min_Rep2, SC_20min_Rep2 overlapped with each other, while the heat shock sample SC_HeatShock_Rep2 diverged into two separate branches indicative of two distinct pseudo-states. When marked by each cell’s cell-cycle phase (Figure 2G), we note that S phase cells dominated the heat-shock sample, as before; both branches of the trajectory taken by the heat shock sample are dominated by S phase cells. On the other hand, the G1, S and G2M phases largely overlap for the control samples, as seen previously. Next we compared the cell-type clustering results with trajectory analysis. Figure 2H shows the pseudo-time trajectory where each cell is colored according to the unsupervised clusters in Figure S3A. The UMAPs in Figure S3B show the contribution of each sample to the overall clustering: we note that cells in cluster 0 come mostly from the control samples (left, middle) and cluster 2 is composed of cells primarily from the heat-shocked sample (right). These results are consistent with Figure S3D where cells in cluster 0 predominantly occur in control samples (left, middle) and cluster 2 in the heat-shock sample (right). DE analysis was performed on the two branches of the heat-shocked sample using Wilcoxon Rank Sum test in Seurat. The expression level in the logarithmic scale was visualized using color gradient on the trajectory tree plot (Figure S3E). Heat shock genes such as *HSC82, HSP12* and *HSP82* (Figure S3E, left, middle, right) are differentially expressed between the two branches of the trajectory indicating differential response in S. cerevisiae cells to heat shock.

### Characterizing expression heterogeneity in *Candida albicans* using mDrop-seq

To demonstrate the utility of mDrop-seq in profiling different yeast species, we applied mDrop-seq to *Candida albicans. C. albicans* is a common hospital-acquired infection that can be life-threatening.[28] This yeast has several features that make it challenging for droplet single-cell profiling. Thicker cell walls *in C. albicans* compared to *S. cerevisiae* (∼150 nm and ∼120 nm, respectively [22]) make lysis of the *C. albicans* cell wall more difficult. This species also has a hyphal phenotype that can clog microfluidic channels and disrupt flow and droplet generation. The Droplet Yeast Lysis Buffer or DYLB (see **Methods**) used to lyse *S. cerevisiae* was insufficient for *C. albicans* (as assessed under a microscope). The *C. albicans* Lysis Buffer used in our experiments is similar to DYLB but with higher concentrations of both detergent and enzyme (see **Methods**). We performed two replicates of lysis experiments (15, 20, and 25 minutes) to establish the ideal incubation time for lysis and RNA capture of *C. albicans* after droplet encapsulation. The 25 minute lysis from the second replicate did not produce a library due to technical error. Average counts varied between the five libraries (Table S1), consistent with lysis experiments performed in bulk. Gene cutoffs were determined for each lysis time and replicate; this ranged from 175 to 370. A total of 14,320 *C. albicans* cells were detected across datasets. Lysis time of 20 minutes outperformed the other lysis times in both replicates for gene and UMI capture (Table S1). We combined the 15 and 20-min incubation datasets from replicate 1 (CA_20min_Rep1 and CA_25min_Rep1) to construct a combined dataset with 4,006 cells, with lower and upper cutoffs of 220 and 1600 genes per cell, shown in Figure S6. The 15-min lysis experiment yielded poor data that may be attributed to incomplete cell lysis, and was excluded from further analysis.

In Replicate 2, 15 and 20-min libraries (CA_15min_Rep2 and CA_20min_Rep2) were also combined (Figure 3). The gene with highest expression, on average, is a white-phase yeast transcript *WH11* (Figure 3A). *C. albicans* in white phase is the generic, round yeast form, and expected when grown in rich medium [29], as opposed to the elongated, mating competent opaque phase.[30] These morphological forms favor growth unlike the *C. albicans’* filamentous hyphal form.[29] *TDH3*, a gene involved in glycolysis (that also showed the highest expression in *S. cerevisiae* data), is the fourth highest expressed gene in the *C. albicans* dataset in both replicates. Potentially due to the difference in genes and UMI detected, we see separation in PC space between CA_15min_Rep2 and CA_20min_Rep2 datasets in Figure 3D (Pearson correlation = 0.96), necessitating batch correction. We see several distinct clusters (clusters 2, 3, and 5) after batch correction in Figure 3E that separate out from the rest of the cells. A heatmap of the most significantly expressed genes in each cluster is shown in Figure 3I. The upper cluster 3 is marked by the expression of GPI-anchored cell wall genes such as *FGR41* and *PGA38* (Figure 3F).

**Figure 3:**
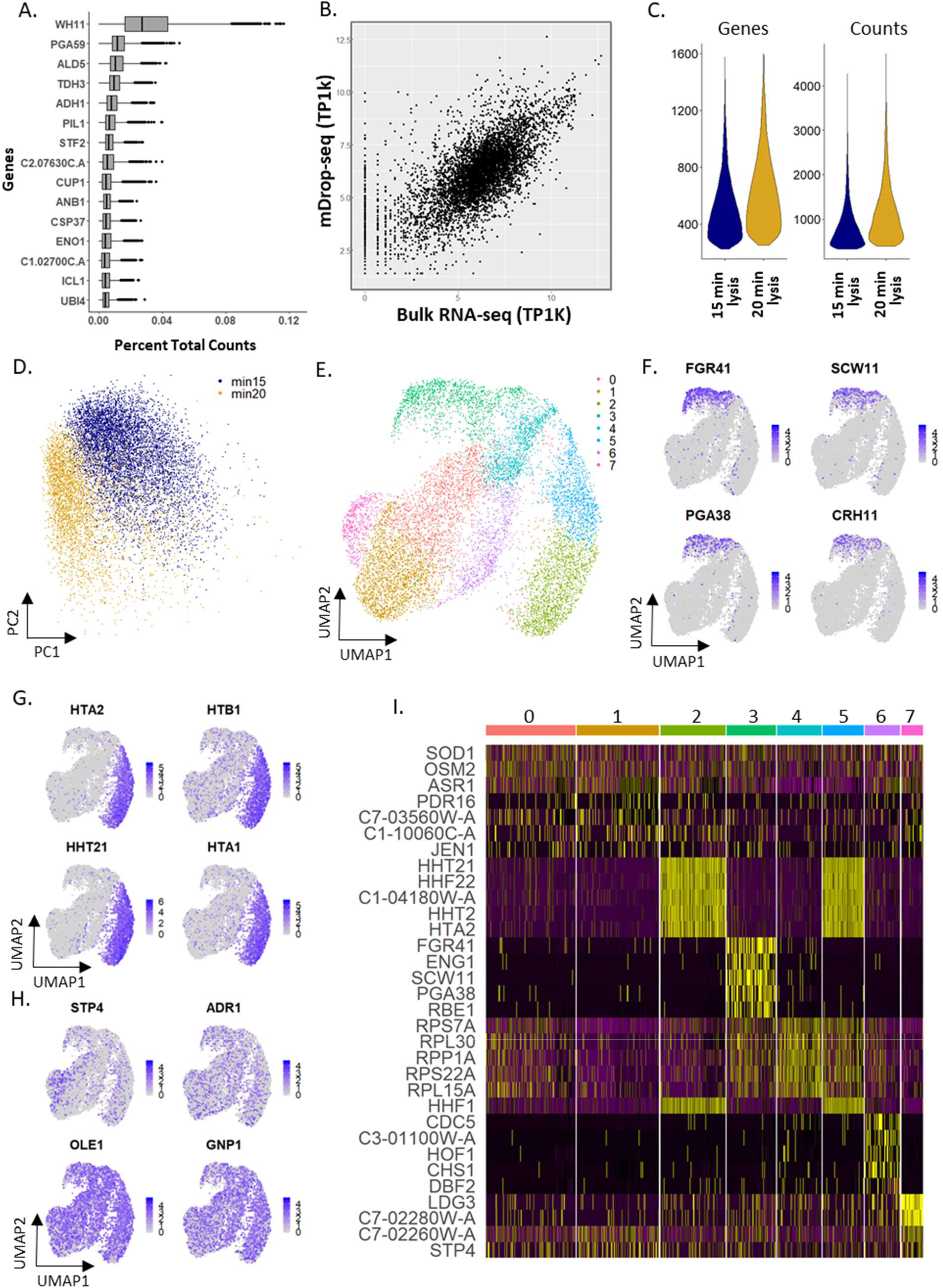
mDrop-seq of 10,314 *Candida albicans* cells. A) A boxplot of 15 genes with highest expression in *C. albicans*, plotted by percentage of total counts. B) Plot comparing average gene expression between CA_rep2 data and bulk RNA-seq. C) Violin plots displaying the genes and UMI counts per cell. D) Plot of PC 1 and 2 in mDrop-seq data for 15 and 20 min incubation times. E) UMAP displaying the clustering analysis of 10,314 *C. albicans* cells detected after batch correction. F) Feature plots displaying 4 GPI-anchored cell wall proteins that represent markers for cluster 3. G) Feature plots displaying 4 histone tail genes that represent markers for clusters 2 and 5. H) Feature plots of transcription factors, fatty acid biosynthesis, and hyphal formation genes involved with *C. albicans* virulence. I) Heatmap displaying expressions of the top marker genes for each cluster.

Clusters 2 and 5 are heavily represented by histone tail genes as markers (Figure 3G), indicating that these clusters may represent variation caused by the cell cycle. Transcription factors *STP4* and *ADR1* encoding for zinc finger proteins and implicated in *C. albicans* virulence [31] and *GNP1*, a transmembrane transporter of amino acids are moderately expressed across the entire population (Figure 3H). *OLE1*, involved in filamentation, shows high expression across all cells, shown in Figure 3H. Figure S5 shows UMAP plots of samples CA_15min_Rep2 and CA_20min_Rep2 without batch correction, where each cell is colored by unsupervised clusters, sample of origin and cell cycle phase in Figures S5A, B, C respectively. Figures S5D, E plot the expression levels of histone tail genes and GPI-anchored cell wall genes in this dataset.

To compare mDrop-seq of *C. albicans* with bulk RNA-seq, we performed a population-level RNA-seq experiment on *C. albicans*. Our population level RNA-seq data showed a modest Pearson correlation of 0.70 (Figure 3B) with pseudo-bulk [3] data constructed from mDrop-seq and 0.64 (Figure S6B) with a publicly available dataset of *C. albicans*.[27] The correlation between our population level data and the public dataset was 0.84, by comparison.

### Differential expression in *Candida albicans* in response to fluconazole exposure

Fluconazole is an anti-fungal agent commonly used to treat infections from various *Candida* species. *C. albicans* cells were exposed to fluconazole over the course of 3 hours, with samples taken for running mDrop-seq prior to exposure, at 1.5 hr and 3 hr. Fluconazole is a bis-triazole antifungal agent that binds to cytochrome P-450 to disrupt the conversion of lanosterol to ergosterol.[32] Previous experiments showed that *C. albicans* responds to the presence of fluconazole in a variety of ways, such as increasing expression of the drug target genes, increasing drug efflux, and finding compensatory pathways for ergosterol biosynthesis.[33] We sampled a population of *C. albicans* cells before (as control) and after (1.5 and 3 hr) exposure to 15 µg/mL fluconazole, which is slightly higher than the C_max_ dose of 400 mg.[34] These experiments were performed twice.

We saw an increase in UMI and genes detected per cell when exposed to fluconazole, across replicates (Figure 4A). Figure 4B shows a UMAP plot of the integrated dataset (control, 1.5 and 3 hr) from Replicate 1 with slight separation between the control and fluconazole-exposed samples. In contrast, there was very little separation between the samples at 1.5 and 3 hour fluconazole exposures. The mean gene expression of the control library yielded a Pearson correlation of 0.91 for the 1.5-hr and 0.88 with the 3-hr time points (Figure 4C).

**Figure 4:**
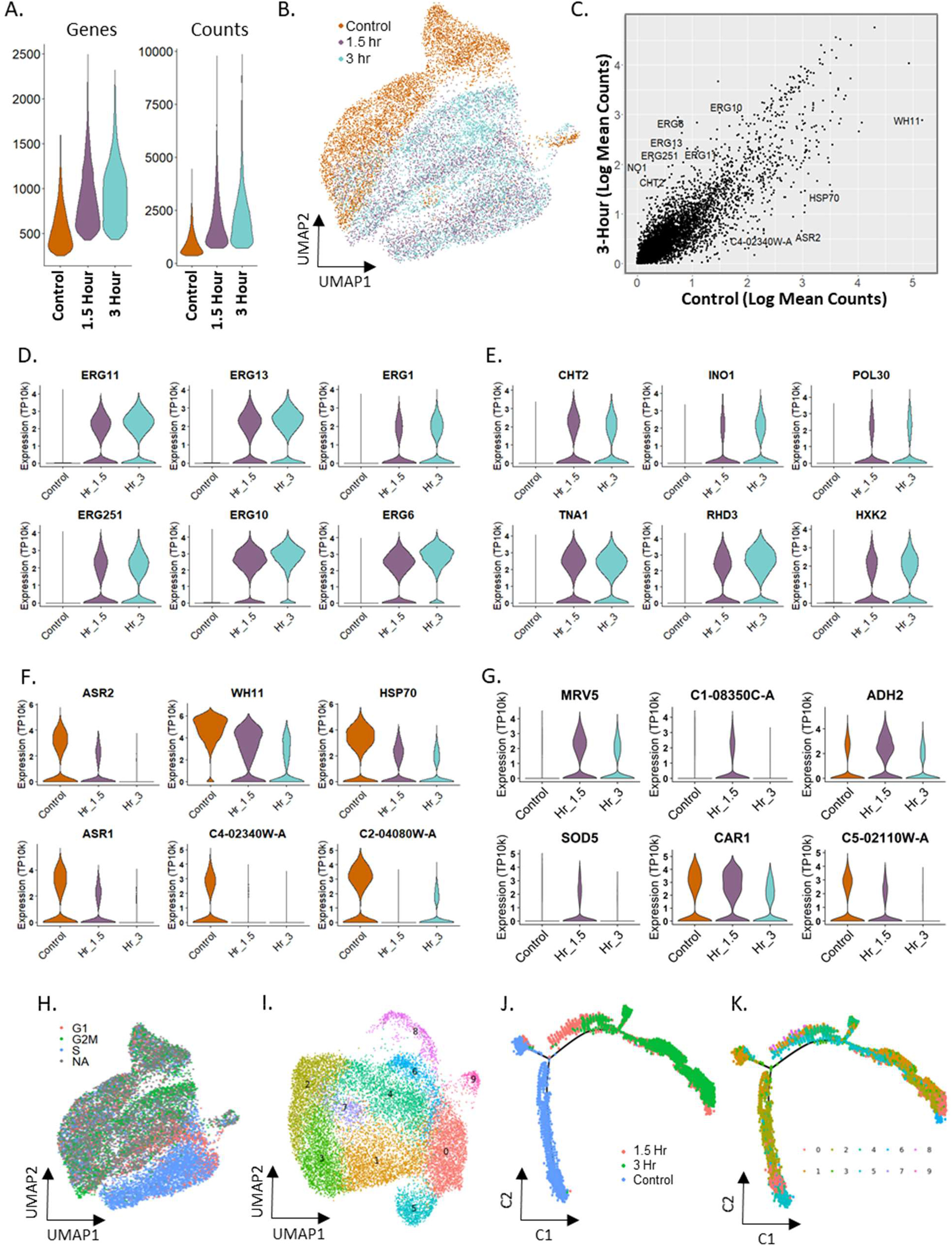
Fluconazole treatment of 15,503 *Candida albicans* cells profiled using mDrop-seq. A) Violin plot displaying the number of genes and UMI for each fluconazole time point B) UMAP displaying the clustering patterns of the integrated control and fluconazole libraries. C) Correlation between the control and 3-Hour fluconazole exposed yeast for gene expression. D) Violin plots displaying the expression differences in control and fluconazole datasets for several ergosterol biosynthesis genes. E) Violin plots displaying the expression differences of genes detected at significantly higher expression in fluconazole treated data. F) Violin plots of genes that show significantly higher expression in the control data. G) Violin plots of DEGs for the 1.5 hour fluconazole exposed data. H) Cell Cycle phase assignment on the integrated UMAP. I) UMAP displaying the integrated data after cell cycle regression. J-K) Pseudo-time trajectory of the combined dataset inferred using Monocle. Colors indicate J) experimental time points, and K) cell-type clusters shown in I.

When comparing the 1.5 and 3 hr time points of fluconazole treatment to the control, we saw significant upregulation of several ergosterol biosynthesis pathway genes that alter *C. albicans* susceptibility to different classes of antifungal drugs like azoles and allylamines.[35] Figure 4D shows six of these genes, with *ERG11* being the main drug target of fluconazole and *ERG1*, associated with terbinafine resistance [36] (LogFC = 1.26, 1.71, 1.02, 1.53, 1.34, and 2.00 for *ERG11, ERG252, ERG1, ERG13, ERG10*, and *ERG6*, respectively). ABC transporters used for drug efflux, were not detected, likely due to the short duration of fluconazole treatment.[33] The fluconazole treated cells also showed increased expression of many histone genes (LogFC > 2) compared to control (Figure S10A). Note that histone genes in yeasts have poly-adenylated tails, unlike in humans.[37]

Using DE analysis on the combined dataset, we identified the following genes of interest: DE genes that show higher expression in the fluconazole treated datasets, e.g., *CHT2, INO1, POL30, TNA1, RHD3, HXK2* (Figure 4E) including several antigenic genes and genes upregulated during a host immune response; DE genes that show increased expression in the control data that decreased with time under fluconazole treatment, e.g., *ASR1, ASR2, WH11, HSP70, AHP1* (Figure 4F) associated with core and heat-shock specific stress responses; and DE genes that show highest expression transiently in the 1.5 hr fluconazole treated sample, e.g., *MRV5, ADH2, SOD5, CAR1* (Figure 4G) associated with acid, osmotic and alkaline stress responses. Violin plots of housekeeping genes *ACT1, PDA1, TDH3*, and *PGK1* are shown in Figure S11B for comparison.

Next, we performed cell cycle analysis on the combined dataset (CA_Fluconazole-ctrl, 1.5, 3hr_Rep1) from the control, 1.5 and 3 hr time points of the fluconazole treatment. The fluconazole treated datasets showed separation of S phase cells from the rest of the cells in PC space (Figure S10B). When colored by cell-cycle phase, the UMAP of the combined dataset (Figure 4H) showed some cell clustering by their assigned cell cycle phase. We also saw significant increase in the number of cells assigned to the S phase under fluconazole treatment (3.2x and 1.8x for 1.5 and 3 hr, respectively; Supplementary Table S2) with respect to the control dataset. Cells under stress tend to go into cell cycle arrest.[38] Increased expression of ERG genes has also been associated with slow growth in yeasts.[39] Since many histone genes occur in the list of marker genes for the S phase, we verified that high histone activity alone in the fluconazole-treated cells was not skewing our cell cycle assignment towards the S phase (see **Analysis of cell cycle phases in *C. albicans*** in Supporting Material).

Unsupervised clustering of the combined CA_Fluconazole_Rep1 data after cell cycle effects were regressed out (Figure 4I) showed clusters of cells exhibiting histone activity (cluster 4), ribosome activity (clusters 4, 6), synthesis of ribonucleoproteins and Hap43 induced proteins, along with reduction in iron metabolism (cluster 5; violin plots of expression for some genes in this cluster are shown in Figure S11C), stress response (cluster 7), synthesis of cell wall and vacuolar proteins (cluster 8), and nucleolar activity (cluster 9). When cells from each time-point were plotted separately (Figure S11A), we saw that some cell clusters were present predominantly in either control or fluconazole-treated time-points, e.g., clusters 5 and 7 in control, cluster 4 at 1.5 hr, and cluster 6 at 3 hr. The number of cells in clusters 2 and 3 decreased monotonically between control, 1.5 and 3 hr time-points. Similar analyses of fluconazole treated *C. albicans*, replicate 2 are shown in Figures S9 and S12A, B. These results show interesting variability in transcriptomic response between cells to fluconazole, potentially providing insight into differences in resistance between cells.

### Trajectory inference in fluconazole stimulation in *Candida albicans*

Since the fluconazole treatment led to steady changes in gene expression along the 3 hr time course, we attempted to capture the temporal changes in gene expression by constructing pseudo-time trajectories for the *C. albicans* stimulation, using Monocle, an R package.[27] We assumed the control (untreated) sample as time t=0 hr in the fluconazole treatment for this analysis. Figures 4J, K show the pseudo-temporal trajectories of CA_Fluconazole_Rep1 response to fluconazole, marked by experimental time-point and cell-type clusters (identified by unsupervised clustering and shown in Figure 4I), respectively. Based on prior knowledge, the tip (bottom left) in Figure 4J was set as the starting point for pseudo-time construction. Occurrence of the untreated control (blue) on the left of trajectory in Figure 4J and the fluconazole treated samples (coral-1.5 hr; green-3hr) to the right are consistent with the pseudo-time progression shown in Figure S11D, as inferred by Monocle. Figure S11E shows the increasing expression of *ERG10* and *ERG11* genes that mediate resistance to fluconazole and other antifungal agents along the trajectory.[38]

The contributions of each experimental time point to the different branches of the pseudo-time trajectory are shown in Figures S11F, G broken down by control (left), 1.5 hr (middle) and 3 hr (right) time points and marked by cell-type clusters and cell cycle stages, respectively. The branches to the left were predominantly composed of cells from the control sample. S phase assignment dominated the fluconazole treated cells (Figure S11G, middle, right), as seen from cell clustering.

Similar analyses of fluconazole treated CA, replicate 2 (CA_Fluconazole_Rep2) are shown in Figures S9 and S12. Again, the representation of each sample along the pseudo-time trajectory is consistent with experimental time points (Figure S9J), starting with control (left) cells at the tip and followed by 1.5 hr fluconazole-treated cells (middle) and 3 hr treated cells (right). Cells belonging to the S phase also dominate the fluconazole treated samples (Figure S12E, middle, right), as seen in replicate 1.

In summary, we show that pseudo-time analysis of CA exposed to fluconazole show that cell activation trajectory in pseudo-time can be used to infer the temporal sequence of gene expression in yeast cells under external stimuli.

## Discussion

As noted earlier, single cell genomic analyses of microbial species have been difficult due to challenges in single cell lysis and low input material in microbial cells. Yeasts and other fungi have poly-adenylated tails on the 3’ end of their mRNA, allowing selective mRNA capture using poly-dT oligonucleotides that is not possible in bacterial cells, making fungi more experimentally tractable among microbial species.

We established the feasibility of mDrop-seq to profile transcriptional heterogeneity in fungal species at single cell resolution and at scale by performing mDrop-seq on a total of 35,109 single cells of *S. cerevisiae* and 39,705 *C. albicans* cells across multiple replicates, experimental conditions and environmental stimuli in the form of heat-shock (*S. cerevisiae*) and fluconazole exposure (*C. albicans*). Based on Drop-seq and DroNc-seq [1],[21] used to profile gene expression in mammalian cells, mDrop-seq leverages existing single-cell experimental and computational tools and allows for lower barrier of entry and easy adaption of single cell RNA-seq on fungal species for labs that are set up for Drop-seq or similar workflows.

### Droplet content and stability

To implement mDrop-seq, we needed to overcome the challenge of microbial cell lysis in emulsion drops, while maintaining droplet stability and RNA integrity for downstream molecular biology reactions. This was accomplished by using a combination of zymolyase and Sarkosyl activity in drops, along with thermal incubation. The enzyme, zymolyase targets a common component of fungal cell walls and requires thermal activity; the detergent, Sarkosyl is a strong lytic agent that works ubiquitously on mammalian cells, zebrafish, *C. elegans* and fruit fly.[40]–[42] Different concentrations of zymolyase and Sarkosyl were used in the mDrop-seq lysis buffers for *S. cerevisiae* and *C. albicans* (see **Methods**), based on bulk and droplet-based lysis experiments on the two species. We posit that similar cocktails consisting of zymolyase and Sarkosyl will prove effective on a broad class of fungal species that share similar cell wall properties to *S. cerevisiae* and *C. albicans*, including clinically relevant species like *C. auris*. Anti-fungal peptides [43] that target specific components of the fungal cell wall may also be added to the lysis cocktail of zymolyase and Sarkosyl.

The stability of emulsion drops and efficacy of downstream reactions are affected by droplet contents. We note that a high concentration of detergent in the lysis buffers, e.g., 3.3% Sarkosyl in the *C. albicans* lysis buffer, is detrimental to stable droplet formation, necessitating lower flow rates on the microfluidic device. Since reverse transcription in mDrop-seq was performed outside microfluidic drops after the emulsion was broken following single cell lysis and mRNA capture on barcode beads in drops [1], [21], the compatibility of lysis buffer and reverse transcriptase was not an issue. The DYLB and *C. albicans* lysis buffer used in our experiments were optimized for the microfluidic device [21], oil-surfactant mix, and flow parameters used here. While stable droplets may be generated at higher flow-rates with other surfactants, we prefer the oil-surfactant mix used here due to its relatively low cost, long shelf-life and easy availability.

### Sequencing and alignment

We used Illumina Paired End (PE) sequencing for mDrop-seq. Due to the relatively lower complexity of the yeast mDrop-seq libraries in general, or GC content bias (∼37%, in *C. albicans* and (∼41% in *S. cerevisiae*, reported by *FastQC*), we found it beneficial to sequence these libraries multiplexed with more complex libraries like those from human (∼45%, from *FastQC*) or use higher Illumina PhiX concentration to improve the overall quality of sequencing runs.

The 20 bp long Read1 sequence was used to de-multiplex the cell barcode and UMI while the 60 bp Read2 was used to identify the 3’ end of transcripts. While longer read lengths can help reduce multi-mapping in complex genomes such as the human at 3.1 Bb [44], yeast genomes are typically much smaller, e.g., 12 Mb for *S. Cerevisiae* [45] and 14.7 Mb for *C. albicans* [46]; transcripts with shorter read lengths (∼30 bp) can be uniquely mapped to them. Figure S7A shows the percent of uniquely mapped reads (left) and reads mapping to multiple loci (right) as functions of Read2 fragment lengths for *S. cerevisiae, C. albicans* and human genomes. We also compared the effect of clipping the transcript fragments on the 3’ vs. 5’ ends on STAR [47] aligner and saw no noticeable difference in mapping rates between the two.

UMI identifies individual mRNA molecules, allowing us to collapse PCR replicates and prevent PCR bias. Across all mDrop-seq experiments, we saw ∼30-120% more UMI compared to the number of genes detected, with higher percrent UMI’s detected in the heat-shock experiments in *S cerevisiae* or fluconazole treatment in *C albicans*, compared to their respective controls. The majority of genes being detected had only a single count of transcript attributed to them. This is expected because most yeast genes are expressed as single copies of mRNA at any time.[23] This may make it difficult to differentiate between true variation and drop-outs in the data, a problem in scRNA-seq that is exacerbated in yeasts.

### Analysis of stress, heat shock and response to anti-fungal agent in yeast cells

When comparing the control and heat shock sample in *S. cerevisiae*, we see a very clear heat shock response and notable separation in PC and UMAP space. This separation is primarily driven by upregulation of known heat shock genes along with genes involved in DNA replication stress (e.g., *KAR2, LST8, ERO1)* and protein transport (e.g., *RTC3, NCE103, TMA10*). Using a pseudo-time trajectory analysis, we clearly see progression from a non-stimulated control to the heat stimulated cells, with a branch point indicating two separate heat-shock responses. The most significant difference between these two branches is the differential expression of ribosomal structure (e.g. *RPL8B, RPL25, RPL36B*) vs. oxidative-reduction energetic processes (e.g. *PIG2, PCL8, GDB1*).

The *S. cerevisiae* data were normalized and batch-corrected to allow comparisons between experimental conditions, and categorized into three sub-groups: control, “stressed” control, and heat-shock, for comparison. We find the “stressed” control group to be intermediate between the control and heat-shock in that it showed an elevated stress-response signature for *HSP12, HSP26* and *HSP42* compared to the control set, but lower than the heat-shock data. In addition, the heat shock data showed expression of genes related to heat-shock stress, e.g., *HSP30*, and *PIN3*, DNA replication stress, e.g., *LST8, BTN2, ERO1*, and protein transport, e.g., *RTC3, TMA10* and *SPI1* that were absent in the control and “stressed” control samples. We saw similar levels of transcription for housekeeping genes, e.g., *ACT1, HTB1, TDH2*, and *FBA1* across all groups. Also of interest are genes like *AIM44, PIR1, PST1*, and *EGT2*, associated with cell wall stability and cell budding that were expressed in a subset of cells clustering together in the control data.

Using gene lists specific to the G1, S and G2M phases to the cell-cycle, we scored and assigned each cell to a unique cell-cycle phase for both *S. cerevisiae* and *C. albicans*. Cells that could not be unambiguously assigned to any particular cell-cycle phase were marked as NA. In both heat shock and drug treatment experiments, we see significant decrease in the number of G2M phase cells (Supplementary Table S2) and an increase cell numbers assigned to the S phase. Since fluconazole treatment may be expected to elicit a stress response in yeasts, we propose that yeast cells are in general likely to get arrested in the S phase under stress.

Since fungal histone mRNA are polyadenylated [37] and thus amenable to mDropseq capture, we were able to characterize histone activity in *S. cerevisiae* and *C. albicans*. Overall, we saw significant differential expression of genes associated with histone activity in the cellular sub-sets identified by unsupervised clustering of both *S. cerevisiae* and *C. albicans* populations. The increase in the number of cells expressing histone genes in environmentally perturbed datasets may be due to stalling of the cell cycle in the S phase. The increase in the fraction of the cells belonging to the S phase may also be due to higher chromatin accessibility needed for increased transcription under stress response that drive up histone expression.

mDrop-seq on *C. albicans* cells exposed to the anti-fungal agent fluconazole showed significant increases in the number of counts and genes detected, compared to control data. Much of these increases appeared to be driven by increased expression of ribosomal structure genes (e.g., *RPS27, RPS6A, RPS12*). Across both replicates, we noted significant up-regulation of several genes belonging to the Ergosterol Biosynthesis pathway, including *ERG11* that produces the cytochrome P-450 target of fluconazole. The upregulation of this pathway is a known mechanism for resisting the membrane disruptive effects of fluconazole. We did not detect any upregulation of ABC transporter genes as mechanisms of resistance. During the 3 hr. exposure time, we also noted several genes that increased their expression transiently, increasing quickly in expression in 1.5 hr. before decreasing by the end of 3 hr. period. These genes included some stress-induced genes such as *SOD5* and *ADH2*, as well as *C1-08350C-A, C5-02110W-A*.

After integrating the control and fluconazole treated data for each replicate, followed by cell cycle regression and clustering analysis, we noted that the signatures of ribosomal and histone expression differences persisted within the clusters. A cluster marked by GPI-anchored proteins was also seen in the integrated data, as well as a cluster involving nucleolar and pre-ribosomal genes.

In particular, we noted increased histone activity in *C. albicans* under fluconazole exposure. Since many cell-cycle genes for the S phase in *C. albicans* are histone-related, we confirmed that the signal for the S phase assignment was preserved, compared to the G1 and G2M phases even when the histone genes were excluded from the S phase marker list (**<u>Supplementary Material</u>**). Since chromatin accessibility is needed for increased transcription under stress response, this may drive up histone expression under heat shock in *S. cerevisiae* or fluconazole treatment in *C. albicans*.

### Scalability as a technology

mDrop-seq offers cheap transcriptomic profiling solution for unicellular fungi and may be easily adapted in laboratories that use Drop-seq or similar techniques. Existing bioinformatics and statistical tools for single cell analyses may be effectively leveraged to analyze fungal species at single cell resolution. According to our estimate, the cost of single-cell library preparation using mDrop-seq is ∼$250 per sample (∼5,000 cells/sample). At ∼50 million reads per sample, we estimate sequencing to cost an additional ∼$190 per experiment.

## Conclusion

We introduce mDrop-seq, a droplet based RNA sequencing of fungal species at single cell resolution and high throughput. We applied mDrop-seq on two important yeast species, viz., *S. cerevisiae*, a model organism commonly used to study fundamental processes in biology, and *C. albicans*, a common, clinically relevant pathogen. We were able to identify cellular subsets and their expression profiles within the larger population that show cell-budding activity, histone synthesis, etc. To our knowledge, this is the first work that performs single-cell RNA-seq on *C. albicans* at high throughput. Modestly priced and based on established protocols that are easy to implement, we anticipate that mDrop-seq will be instrumental for understanding the basis on phenotypic and functional variability in fungal species in a wide range of contexts, including medical treatment of fungal pathogens, understanding basic biological processes in model organisms, and production of biologics.

## Acknowledgment

We thank Sean Crosson for help setting up *C. albicans* cell culture, Ashish Thakur for help designing bulk RNA-seq experiment, Samantha Keyport for help designing heat shock experiments and Aviv Regev, Carl de Boer and Sebastian Pott for discussions and feedback. This work was supported by NIH R21 AI144417-02, DP2 AI158157-01 and BSF 2017357 grants. RPR was supported by NIH U19AI110818 and 1R15AT009926-01. Next-Gen sequencing was performed at the University of Chicago Functional Genomics Facility and computational resources were provided by the University of Chicago Research Computing Center.

## Materials and Methods

### Yeast strains and cell culture

*Saccharomyces cerevisiae* (strain BY4741, Open Biosystems) cells were grown as a dense culture in YEPD (MP Biochemicals, #: MP114001022) at 27 °C overnight. The *S. cerevisiae* culture was heavily diluted (1:20) in fresh YEPD medium and grown for 4 hours at 27 °C, following which the cells were placed on ice and chilled. *Candida albicans* (strain SC5314, ATCC) were grown in YEPD at 27 °C after heavy dilution (∼1:375). After 20 hours of cell culture, *Candida albicans* cells were chilled on ice prior to processing using mDrop-seq.

### Heat shock stimulation of *S. cerevisiae*

After *S. cerevisiae* was grown in YEPD for 4 hours post dilution, cells were counted in YEPD using Neubauer Improved (NI) hemocytometer (InCyto, #DHC-N01-2). 700,000 cells were aliquoted into a 1.7 mL microfuge tube and heat-shock stimulation was applied by placing the tube in an Eppendorf F1.5 Thermomixer set to 42 °C and 500 rpm for 20 minutes. At the end of the heat shock incubation, the microfuge tube containing the *S. cerevisiae* cells were placed on ice for 5 minutes. The cells were then washed once with ice-cold 1X PBS (Teknova, #P0195) and 0.01% BSA (NEB, #B9000Sm), henceforth referred to as PBS-BSA, quickly recounted, and brought to a concentration of 700,000 cells/mL in PBS-BSA. 10 µL of RNase Inhibitor was added to 1 mL of yeast cells in PBS-BSA and mDrop-seq was performed as described below. The emulsion droplets were collected on ice to preserve the heat shock signal during the droplet encapsulation period.

### Fluconazole stimulation of *Candida albicans*

After *C. albicans* was grown overnight in YEPD, 5 million cells were counted using a Neubauer Improved (NI) hemocytometer (InCyto, #DHC-N01-2) and diluted into 2 mL of fresh YEPD. 1 million cells were removed from this pool and put on ice as the control population and processed using mDrop-seq. Fluconazole (Sigma, #F8929-100MG) was freshly diluted to 100 µg/mL in fresh YEPD, and added to the remaining 4 million *C. albicans* to a final concentration of 15 µg/mL. The *C. albicans* was then incubated in fluconazole under end-over-end rotation at room temperature for 1.5 and 3 hr, removed and put on ice prior to running mDrop-seq.

### mDrop-seq cell preparation and co-encapsulation in droplets

Yeast cells (*S. cerevisiae* or *C. albicans*) were centrifuged separately at 1000 xg for 3 minutes in a swinging bucket centrifuge at 4 °C. The cells were washed twice with ice cold PBS-BSA. Following the washes, 10 µL of cells was sampled and counted using a NI hemocytometer (InCyto, #DHC-N01-2). ∼1 mL of cells at 700,000 cells/mL suspended in PBS-BSA was placed in a 2.5 mL syringe (BD, #309657). 10 µL of RNase Inhibitor (Lucigen, #F83923) was added per 1 mL suspension immediately before microfluidic encapsulation. A 75 μm DroNc-seq device was used for droplet generation.[21] Cells and beads were co-flowed into the microfluidic device, both at 1.5 mL/hr for *S. cerevisiae* and 1 mL/hr for *C. albicans*. Cells at 700,000 cells/mL and 4,500,000 droplets/mL gives a Poisson loading distribution with λ = 0.15.

Barcoded beads (ChemGenes, #Macosko-2011-10(V+)) were suspended in DYLB or *C. albicans* lysis buffers at 350,000 beads/mL and kept in suspension by constant stirring with a magnetic tumble stirrer and flea-magnet setup (V&P Scientific, #VP 710, #772DP-N42-5-2); the flea magnet is placed in the syringe containing the barcode beads suspended in lysis buffer and the stirrer kept in the vicinity of the syringe during droplet generation. Cells and beads in lysis buffer were co-encapsulated in drops using a surfactant-oil mix (BioRad, #1864006) flowed at 8 mL/hr in a 10 mL syringe (BD, #302995) as the outer carrier oil phase. Droplets were collected at ∼3750 droplets/sec for 30 minutes in 50 mL tubes (Genesee Scientific, #28-106).

*Saccharomyces cerevisiae*: Barcoded beads were suspended in Droplet Yeast Lysis Buffer or DYLB, consisting of 6.7% (w/v) Ficoll PM-400 (GE Healthcare, #17-0300-05), 225 mM Tris pH 7.5 (Teknova, #T2075), 22 mM EDTA (Fisher, #BP2482-500), 0.67% Sarkosyl (Teknova, #S3377), 55 mM KH_3_PO_4_ (Sigma, #P5629-25G), 1.3 mM DTT (Teknova, #D9750), 0.1% (v/v) β-mercaptoethanol (Sigma, #M6250-10ML), and 450 units/mL zymolyase (Zymo Research, #E1005); this mix was optimized for *S. cerevisiae* lysis in drops.

*Candida albicans*: Barcode beads were suspended in *C. albicans* lysis buffer containing 6.7% Ficoll PM-400 (GE Healthcare, #17-0300-05), 225 mM Tris pH 7.5 (Teknova, #T2075), 22 mM EDTA (Fisher BP2482-500), 3.3% (w/v) Sarkosyl (Teknova, #S3377), 55 mM KH_3_PO_4_ (Sigma, #P5629-25G), 1.3 mM DTT (Teknova, #D9750), 0.1% (v/v) β-mercaptoethanol (Sigma, #M6250-10ML), 650 units/mL zymolyase (Zymo Research, #E1005), that was optimized for lysing *C. albicans* in drops.

*S. cerevisiae and C. albicans species mixing:* Each yeast cell population, *S. cerevisiae* or *C. albicans* were processed separately as follows: Each yeast species was centrifuged at 1000 xg for 3 minutes in a 4 °C swinging bucket centrifuge and washed twice with ice cold PBS-BSA. Following the washes, 10 µL of each cell aliquot was sampled from each species and counted using a NI hemocytometer.

Two experiments at two different Poisson loading concentrations, were performed to calculate doublet rates at these loading conditions: 175,000 cells from each species were combined and the final volume adjusted to 1 mL of PBS-BSA for λ ≈ 0.077; 350,000 cells from each species were suspended in a total volume of 1 mL PBS-BSA for λ ≈ 0.15. Due to the presence of *C. albicans*, the stronger *C. albicans* lysis buffer was used for both experiments.

### Cell lysis, reverse transcription, cDNA amplification and Nextera library generation for mDrop-seq

After droplet collection, the 50 mL tubes were transferred to a 37 °C water bath for zymolyase digestion and lysis for ∼20 minutes; different lysis incubation times ranging from 10-25 mins were tested, both for *S. cerevisiae* and *C. albicans*. After the incubation, the Drop-seq protocol was followed for breaking droplets, collecting barcode beads with mRNA hybridized onto them and washing them in 6x Saline-Sodium Citrate (Teknova, #S0282).[1] Reverse transcription was performed in 1.5 mL microfuge tubes under end-over-end rotation using a modified Reverse Transcription mix (1x Maxima H-RT buffer, 4% Ficoll PM-400 (GE Healthcare, #17-0300-05), 3 mM MgCl_2_ (Sigma, #M1028), 1 M Betaine (Sigma, #14300), 1 mM dNTPs (Clontech, #639125), 1 U/μL Rnase Inhibitor (Lucigen, #F83923), 2.5 μM Template-Switching Oligo primer, (AAGCAGTGGTATCAACGCAGAGTGAATrGrGrG), and 10 U/μL Maxima H-RT enzyme (ThermoScientific, #EP0751) with a 30-minute incubation at room temperature, followed by a 90-minute incubation at 50 °C. This generated barcoded cDNA affixed to the barcoded beads referred to as Single Transcriptome Attached to MicroParticles or STAMPs. The beads were then washed once in 0.5% SDS (Teknova, #S0288), twice in 0.02% Tween 20 (Teknova, #T0710) both prepared in TE buffer (Teknova, #T0228), and treated with Exonuclease I digestion (Fisher, #M0293L). The total number of STAMPS collected was counted manually under the microscope. cDNA amplification was performed on RNA-DNA conjugates attached to ∼120,000 barcode beads in a 96-well plate (Genesee Scientific, #24-302) loaded at 10,000 STAMPs per well. The STAMPS were amplified for 17 PCR cycles, using Kapa Hifi Hotstart 2x Mastermix (Fisher, #NC0465187) and SMART PCR primer (AAGCAGTGGTATCAACGCAGAGT).[1] Post-PCR cleanup was performed by removing the STAMPs and pooling the supernatant from the wells together into a single 1.7 mL low-retention tube (Genesee Scientific, #22-281LR) along with 0.6X Ampure XP beads (Beckman Coulter, #A63880).[1] After adding the Ampure beads to the PCR product, the tube was incubated at room temperature for 2 minutes on a thermomixer (Eppendorf Thermomixer C, #5382000023) set to 1250 rpm, and for another 2 minutes on bench for stationary incubation. Next, the tube was placed on a magnet, and washed 4X times using 1 mL ethanol (Sigma, #E7023) at 80% concentration in each wash. cDNA was eluted in ultra-pure water (Life Tech, #10977-023) at 2.5 μL/well and the concentration and library size were measured using Qubit 3 fluorometer (Thermo Fisher) and BioAnalyzer High Sensitivity Chip (Agilent, #5067-4626). Representative traces of cDNA libraries of *S. cerevisiae* and *C. albicans* are shown Figure S13A, B respectively.

500 pg of the cDNA library was used in Nextera library preparation, following the original Drop-seq protocol, with a 3-minute 72 °C incubation step added at the beginning of the thermo-cycling program to yield Nextera libraries averaging 500-700 bp. [1], [21]

### Population level RNA-seq library preparation of *Saccharomyces cerevisiae* and *Candida albicans*

We performed the same standardized procedure to prepare bulk RNA-seq libraries for *S. cerevisiae* and *C. albicans* as follows: We lysed ∼8,000,000 cells each using DYLB (*S. cerevisiae*) and *C. albicans* lysis buffer (*C. albicans*) and incubation at 37 °C for 20 mins. Total RNA was extracted from each lysate using the Direct-zol RNA Miniprep Plus kit (Zymo Research, #R2071), assessed RNA quality using Qubit HS RNA Assay (Invitrogen, #Q32852) and diluted to 25 pg/µL. The total RNA library was annealed to 5’-AAGCAGTGGTATCAACGCAGAGTACTTTTTTTTTTTTTTTTTTTTTTTTTTTTTTN-3’ primer (IDT) that allow polyA selection of mRNA and template switching, similar to Drop-seq. Briefly, 11 µL of total RNA library was mixed with 11 µL of 10 µM primer above, 11 µL of 10 mM dNTP (Takara, #639125), 13.75 µL of ultra-pure water and 2.75 µL of RNase Inhibitor (Fisher, #NC1081844), and incubated at 75 °C for 3 mins on a PCR thermocycler. A reverse transcription master-mix consisting of 11 µL of 5X Maxima RT buffer, 2.2 µL of H_2_0, 11 µL of 5 M Betaine (Sigma, #14300-500G), 1.65 µL of 100 mM MgCl_2_ (Sigma, #M1028), 2.2 µL of 50 µM Drop-seq TSO primer (IDT), 1.1 µL of RNase inhibitor (Fisher, #NC1081844) and 2.75 µL of Maxima H-RTase enzyme (Fisher, #FEREP0753) was added immediately after the annealing step. The final RT reaction volume of 45 µL was pipetted several times to mix, centrifuged briefly to spin down the contents and incubated on a PCR thermocycler using the following program: 42 °C for 90 min; 5 cycles*(42 °C for 2 min, 50 °C for 2 min); 70 °C for 15 min, to perform reverse transcription of the polyadenylated mRNA selectively annealed to the primer above. The cDNA was amplified for 12 cycles for *S. cerevisiae*, and 13 cycles for *C. albicans*, using Drop-seq TSO-PCR primer (IDT) and KAPA HiFi HotStart ReadyMix PCR Kit (Fisher, #NC0465187K). Amplified cDNA was quantified using Qubit and BioAnalyzer, followed by Nextera library generation.

### Sequencing

Nextera libraries of samples, including bulk RNA-seq, were loaded at ∼ 15 pM concentration and sequenced on an Illumina NextSeq 500 using the 75 cycle v3 kits for paired-end sequencing. 20 bp were sequenced for Read 1, 60 bp for Read 2 using Custom Read 1 primer, GCCTGTCCGCGGAAGCAGTGGTATCAACGCAGAGTAC, according to protocol. Due to the low complexity of yeast cDNA libraries, Illumina PhiX Control v3 Library was added at 5-10% of the total loading concentration for all sequencing runs. Samples for each experiment were loaded at 7-15% of a NextSeq 500 lane and yielded 10-40 million reads for each sample. Some samples were sequenced twice, depending on library complexity. For these samples, the Fastq files were concatenated using the UNIX `zcat` function before running the *dropRunner* pipeline. Data are available at: https://www.ncbi.nlm.nih.gov/geo/query/acc.cgi?acc=GSE154515

### mDrop-seq data preprocessing, alignment and quality control

There were ∼5000-10,000 cells processed per sample and each library was sequenced at ∼40-90 million reads. We developed a *Snakemake* [48] protocol called *dropRunner* that takes paired-end reads in FASTQ format as input and produces an expression matrix corresponding to the UMI of each gene in each cell. The protocol initially performs *FastQC* [49] to obtain a report of read quality. The reports are later inspected manually to ensure high-quality reads were generated. Next, it creates a whitelist of cell barcodes using *umi_tools* [50] *0*.*5*.*3*, which is a list of the top 10,000 valid cell barcodes in terms of number of reads. Next, reads were aligned to the respective yeast genomes using *STAR* [47] 2.7.0a. *STAR* 2.7 introduced *STARsolo*, a turnkey solution for processing droplet single-cell RNA-seq data built directly into the *STAR* aligner. The whitelist and paired-end reads are used as input for *STARsolo*, which performs alignment, gene UMI counting, and cell-barcode-filtering in one step. *STARsolo* uses a heuristic approach for filtering cells with low or noise-level UMI counts. It does so by constructing a UMI count rank plot for each cell (a knee-plot) and picks a cut-off based on the knee of the curve. The pipeline can be found at GitHub (aselewa/dropseqrunner). The filtered digital expression matrices from *STARsolo* were loaded in Seurat (v3.1.1), an *R* package for downstream single cell transcriptome analyses.

All data from *S. cerevisiae* were aligned to the *Saccharomyces cerevisiae* (SC) reference genome, version sacCer3_s288c (https://www.yeastgenome.org/strain/S288C) obtained from the *Saccharomyces* Genome Database (SGD), along with gene lists for each cell cycle. *C. albicans* data aligned to *Candida albicans* SC5314 (CA) reference genome, version A21-s02-m09-r10 (http://www.candidagenome.org/download/gff/C_albicans_SC5314/Assembly21/).

### Bulk RNA-seq data processing

Two sets of bulk RNA-seq data obtained from *S. cerevisiae* and *C. albicans* were assessed for read quality using *FastQC*, mapped to the respective genomes described above using *STAR* 2.7.0a aligner [47], and RNA counts were generated from the bam files using *FeatureCounts* [51]. Read lengths were down-sampled during alignment using the *STAR* aligner, ‘--clip3pNbases’ and ‘--clip5pNbases’ parameters. Count matrices for the two yeast species were compared to their corresponding mDrop-seq datasets using the Seurat package.

### Clustering cells and generating UMAP

We followed the analysis pipeline recommended by Seurat. Data were normalized and scaled using default commands provided by the Seurat package in R. Seurat was used to calculate the gene dispersion and mean expression to find highly variable genes. This reduces the computational time of PCA compared to using the full set of genes. Highly variable genes were used to calculate the PCs for the yeast mDrop-seq data. An elbow plot displaying the variation explained by each PC was used to determine the number of PCs needed to explain the majority (>90%) of the variation. The top PCs determined in this way were used to perform clustering which was visualized with Uniform Manifold Approximation and Projection (UMAP) [24].

### Calculating doublet rates from species mixing experiments

An mDrop-seq dataset containing a mix of *S. cerevisiae* and *C. albicans* cells was aligned once to the SC genome and again separately, to the CA genome. Cells were removed based on data quality. The 7% Poisson loading experiment had gene cutoffs of 50 for both experiments, while the 15% Poisson loading experiment had cutoffs of 275 and 400 for *S. cerevisiae* and *C. albicans*, respectively. We identify cell barcodes that capture genes from both SC and CA genomes. Due to the similarities between the yeast genomes caused by shared ancestry, there are genes that will map to both species. This was checked for by taking a *C. albicans* dataset and mapping it to the SC genome, and vice versa, to identify the genes common to both species. After removing these common genes from the mixed-species dataset, we identified the cell barcodes that show significant mapping (> 50 genes) to both genomes as true doublets.

### Batch correction

Dataset merging and batch correction were performed using the *Anchor Integration* function in the Seurat package. Datasets were independently normalized and had highly variable genes detected using gene dispersion and mean expression. The datasets were scaled before running *Canonical Correlation Analysis* (CCA) function in Seurat to determine dataset anchors and merge the objects. A new integrated dataset was then created using the detected anchors. This integrated dataset was used for dimensional reduction and clustering analyses.

### Cell cycle analysis

Lists of genes that serve as cell cycle phase markers for G1, S, and G2M phases were obtained from Spellman et al.,[52] for *S. cerevisiae* and the candida genome database for *C. albicans*.[53] Cell cycle assignment was made based on the G1, S, and G2M markers for cell-cycle phases. The *AddModuleScore* function implemented in Seurat was used to calculate cell-cycle score for each phase. This function sampled random genes as a control set; the number of the genes in the control set was determined by the number of markers in the cell-cycle gene lists. Since there were three such lists of markers, the minimum size of cell-cycle marker lists was used. For each cell, the largest module score was selected among the three phases and the corresponding cell cycle phase for the selected module score was assigned to the cell. A threshold on the selected module score was applied to ensure the cell cycle assignment was robust. If all module scores for a cell were below zero, the phase for the cell was left undecided and counted as ‘Not Assigned’ or NA in Supplementary Table S2.

This module score list was used with Seurat to create a column in the object metadata containing the assigned cell-cycle phase. Cell-cycle phase metadata was used to calculate PCs instead of highly variable genes. Cell-cycle variation was regressed out during data scaling and centering. The expression percentage and level for the G1, S, and G2M marker genes were visualized using dot plots where the size of the dots indicates the percentage of cells expressing a marker and the color intensity reflects normalized expression.

### Hierarchical clustering of cells

To investigate variations in single cell expression that are not related to cell division and proliferation, the cell cycle effects were removed when scaling the gene expression per cell. The module scores for each of the G1, S, and G2M phases were regressed out against each gene. PCs and Shared Nearest Neighbor (SNN) graph [54] were constructed from the scaled gene expression matrix.

### Trajectory analysis on single-cell data

To trace the lineage or process of temporal activation in yeast cells in response to external stimuli such as heat shock or fluconazole treatment, the trajectory analysis [27] was performed on the single-cell data from control and stimulated cells. Datasets from different conditions/time-points were integrated using anchor-based integration described above. The R-based pipeline, Monocle 2 (v 2.10) was used to process the data and construct the trajectory.

The genes used to order cells along the pseudo-time trajectory, or ordering genes, were set based on differentially expressed (DE) genes obtained from unsupervised clustering in Seurat. DE genes with q-value < 0.01 were selected as the ordering genes in Monocle. The count matrix was log-transformed after adding one to the counts to eliminate logarithms of zero values. PCA was performed on the normalized count matrix using the ordering genes and the top variable PCs were selected based on the scree plot of variance explained per component. The PCs were reduced into a tree structure by Discriminative Dimensionality Reduction with Trees (DDRTree).[55] The backbone of the tree branches formed the cell trajectory. The root of the trajectory was set as the tip of the tree branch that contained the largest number of cells from the control sample, and each cell was ordered in pseudo-time based on the root. During the PCA and DDRTree dimension reduction phases, we removed any cell cycle effects by specifying the G1, S, and G2M module scores obtained from Seurat as variables to be linearly subtracted from the data to look for changes in gene expression that are independent of cell-cycle effects as response of external stimuli.

