## Supplemental Figures and Text for "mDrop-seq: Massively parallel single-cell RNA-seq of *Saccharomyces cerevisiae* and *Candida albicans*" (from another version of this preprint: version 1, 2021-07-06)

### Optimization of mDrop-seq for *Saccharomyces cerevisiae*

mDrop-seq requires cell lysis after single cells are encapsulated in droplets. We developed targeted in-drop lysis protocols for each species that consist of cocktails of zymolyase, Sarkosyl and incubation at 37 °C. Sarkosyl is a strong detergent used for cell lysis [56]; zymolyase is an enzyme mixture with optimal activity at 37 °C, commonly used to digest yeast cell wall. We note that zymolyase or Sarkosyl alone were insufficient to obtain total lysis of yeast cells in drops. Because droplet stability is influenced by constituent fluids and flow parameters, we optimized the Droplet Yeast Lysis Buffer (DYLB) composition with oil-surfactant mix, droplet size and flow rates for each yeast species. Monodisperse droplets of 75 µm diameter that are stable under flow and thermal incubation were used for mDrop-seq experiments. The scRNA-seq workflow in yeasts in microfluidic drops is summarized in Figure 1A.

### Batch correction in *Saccharomyces cerevisiae* replicates

The control experiment was repeated with 15 and 20 min incubation times for lysis, labeled SC_15min_Rep2 and SC_20min_Rep2, respectively. For both datasets, we obtained slightly lower numbers of cells, and comparable numbers of genes and UMI per cell within the datasets (Table S1 and Figure S2A). Overall, the two lysis times in replicate 2 yielded comparable data with Pearson correlation of 0.98.

The two replicates, SC_XXmin_Rep1 and SC_XXmin_rep2 (Pearson correlation = 0.86), show overlapping but slightly different sets of differentially expressed genes and some differences in their expression profiles. Several genes that are more highly expressed within replicate 2 included *HSP12*, *HSP42*, *DDR2*, and *NCE103* (Figure S2B), many of which are involved in stress response.

Figure S2C shows a PC plot combining the two replicates of two lysis incubation times into a single dataset of 20,548 cells, with clear separation between SC_XXmin_Rep1 and SC_XXmin_Rep2. Figure S2D shows the same PC plot after integrating the four experimental replicates using the *Anchor Integration* function in Seurat. UMAP of the integrated data shows that cells from the two replicate experiments clustered together (Figure S2E). Figure S2F shows the 15 highest expressed genes in this second replicate, with *HSP82*, *HSP12*, and *HOR7* being among the top 4 genes expressed. This, along with SC_20min_Rep2 having higher genes and UMI detected, compared to SC _15min_Rep2, may indicate that cells in this experiment underwent longer lysis incubation or otherwise experienced stress, yielding higher measure in stress response.

Because of the noted increase in stress response in the second replicate, we compared the second "stressed" control replicate to the heat-shock experiments. Figure S4D shows a UMAP combining all lysis and heat-shock datasets together, without any correction applied for batch effects. Though the separation of the two control replicates in Figure S2C is apparent, we did not pursue this combined dataset further due the increasing complexity in correcting for multiple batches. Instead, replicate 2, or the "stressed" control replicate alone was compared to the heat-shock data. Figure S4D shows a marked separation of replicate 2 from the heat-shock data in UMAP space, perhaps indicating that the heightened stress response in replicate 2 is still lower than the stress-response in the heat-shock data. Indeed, when comparing expressions of *HSP12*, *HSP26*, and *SSA4*, Figure S4E shows increased overall expression in replicate 2 or the “stressed” control, though lower than expression in the heat-shock data (logFC = 1.99, 2.04, 2.29, respectively). Housekeeping genes show similar expression across replicate 2 and heat-shock datasets (Figure S4F).

**Analysis of cell cycle phases in *Saccharomyces cerevisiae***

In an unstimulated dataset sampling continuously cycling cells, one may expect to find variation attributable to cycling cells. Using cell-cycle gene lists provided in by Spellman *et al.* [57], we attempted to assign a unique G1, S, or G2/M stage of the cell cycle to each *S. cerevisiae* cell that passed our quality filter. Because of low transcript counts inherent in yeasts, we were able to assign G1, S and G2M phases to only a subset of cells in each experiment (Table S2). The remaining cells were designated as ‘not assigned’ or NA and their numbers and percentages are also reported in Table S2. We performed PC analysis on the cycling genes in the combined dataset of the *S. cerevisiae* replicates (Figure 1D). Cells in the same stage of cell-cycle largely grouped together with some overlap (Figure S1D). Unsupervised clustering was performed on the combined dataset after regressing out cell cycle effects using Seurat *ScaleData*. When visualized in UMAP space in Figure S1E, cell-cycle genes in the S and G2M phases appear enriched in the upper left clusters (clusters 0 and 3 in Figure 1G, respectively) while G1 and additional G2M cells are spread across the lower clusters (clusters 1, 2, and 5 in Figure 1G). Figure S1F shows six histone genes enriched in cells assigned to S phase, as expected. All cell-cycle phases are represented in the upper right cluster (corresponding to cluster 4 in Figure 1D) in the UMAP. Cluster 2 in Figure S1G shows increased expression of stress response genes, *DDR2, NCE103, HSP12,* etc. (also shown in feature plots in Figure S1H), likely induced during cell lysis. Figure S1I shows expression of genes associated with GPI-anchored cell wall proteins such as *AIM44* and *PIR1* that cluster together, independently of the cell cycle.

**Analysis of potentially stressed *Candida albicans* shows differential expression**

An early lysis time experiment replicate of *C. albicans,* CA_20min_Rep1 and CA_25min_Rep1, showed heterogeneity both between and within experimental libraries. In this replicate, the 15 minute lysis incubation time also produced a usable library, but was not included due to differences in total counts per cell with the other two time points (Table S1). Gene cutoffs were determined for both 20- and 25-minute lysis times at 370 and 250 respectively. A total of 4,006 cells were detected across the two lysis times. Lysis time of 20 minutes outperforms the other lysis times, with an average of 735 genes and 1,356 UMI detected. We combined the 20 and 25-min incubation datasets, CA_20min_Rep1 and CA_25min_Rep1 to construct a combined dataset with lower and upper cutoffs of 250 and 2,000 genes per cell (Pearson correlation = 0.95). Much like CA_15min_Rep2 and CA_20min_Rep2 in Figure 3, combining these datasets required batch correction prior to visualization in UMAP space (Figures S6C, D). The top expressed genes in this dataset are shown in Figure S6A.

We saw several distinct clusters (clusters 6-10) in Figure 3SD that separated out from the rest of the cells. Figure S7B shows that these clusters have slightly higher average number of genes (left) and UMI or counts (right) detected. Cluster 10 represents cells producing histones, with many histone genes coming up as the most significant markers for this cluster (p_adj<5.11e-33; Figure S6G). Cluster 6 shows differential expression of genes associated with cell wall stability, e.g., *FGR41*, *PGA38*, and *SWE11* (p_adj<1e-292; Figure S6E). *CHT3*, a chitinase gene, was also expressed in cluster 6. Transcription factors *STP4* and *ADR1* encoding for zinc finger proteins and implicated in *C. albicans* virulence [31] and *GNP1*, a transmembrane transporter of amino acids were moderately expressed across the population. *OLE1*, involved in filamentation, showed high expression across all cells, shown in Figure S6F.

**Analysis of cell cycle phases in *Candida albicans***

Cell cycle marker genes for *C. albicans,* determined through microarray analysis of synchronized cells, were obtained from the Candida Cell-Cycle Database [53]. Cells from the combined *C. albicans* lysis datasets were scored for G1, S and G2M specific markers and assigned to the cell cycle phase with the highest score (Figure S8A for replicate 1, Figure S5F for replicate 2). As seen before in *S. cerevisiae* cells, we were able to unambiguously assign cell cycle phases to a subset of cells in each *C. albicans* mDrop-seq experiment. The remaining cells were designated as NA (Table S2). The number and percentage of cells in the different cell cycle phases in each dataset are summarized in Table S2. When the cells are colored by its assigned cell-cycle phase on UMAP space, we note that the cells marked as NA (gray) are more or less uniformly distributed across the UMAP plot.

In replicate 1, PC analyses show minimal separation between the different phases (Figure S8B), potentially due to many genes being lost during integration of the datasets. Regressing out the contribution from cell cycle genes from this dataset had minimal effect (Figure S8C). Instead, we used the CA_20min_Rep1 dataset alone to analyze the effect of cell cycle regression. UMAP plots of the CA_20min_Rep1 dataset marked by cluster number and cell phase assignment are shown in Figures S8D and S8E, respectively. Figure S8F shows the CA_20min_Rep1 dataset after the variation from cell cycle genes was regressed out. Regressing out any variation in expression due to the cycling genes can be useful when comparing experimental conditions where the effects may be small (e.g., environmental stimuli).

For the experiments where *C. albicans* cells were treated with fluconazole, we noted significant increase in the fraction of cells assigned to the S phase, and decrease in the fraction of cells assigned to the G2M phase, compared to their controls. The fraction of cells in the G1 phase did not change much, by comparison. These observations are consistent for both replicates of fluconazole treatment. This is also qualitatively similar to *S. cerevisiae* cells that underwent heat shock treatment.

To verify if cell assignment to the S phase was not biased by high expression of histone genes, the expression levels for S phase markers were examined with and without the contribution of histone genes. Figures S10C, E show dot plots of expression levels for S phase marker genes for all assigned phases of the cell-cycle (G1, S, G2M and NA) in the fluconazole datasets, CA_Fluconazole_Rep1 and CA_Fluconazole_Rep2. The average expression levels of all S phase genes in the cells assigned to the S phase (on the top row of the dot plot) were higher than the expression in the remaining cells. Even when the histone genes were excluded from the S phase marker list (Figures S10D, F), cells assigned to the S phase showed higher expression than G1 and G2M phases. This indicated to us that high S phase assignment among the *C. albicans* cells was not driven by the high expression of histone genes alone.

### Supplementary Figures

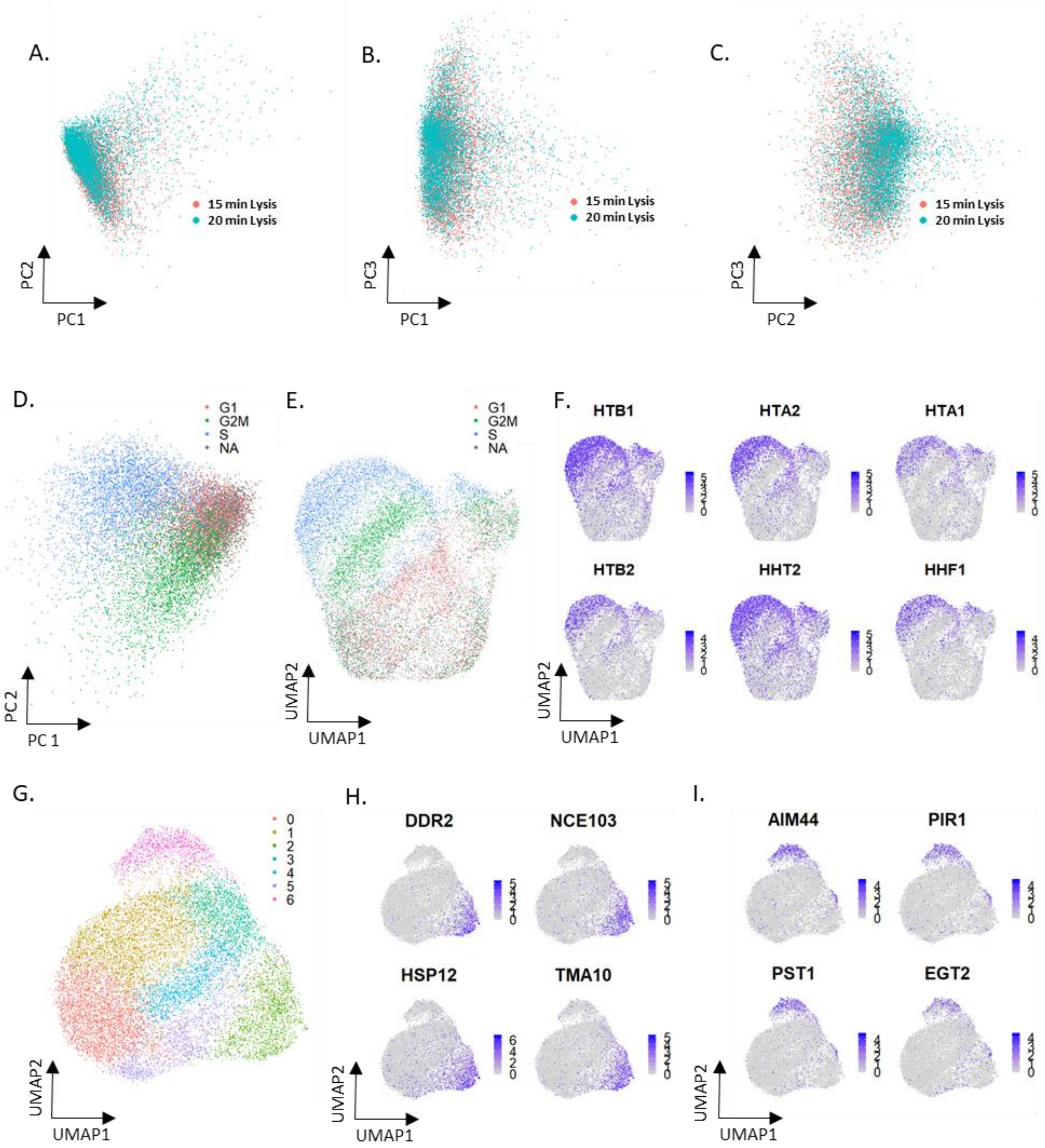


**Figure S1: Analysis of 12,012 *S. cerevisiae* cells, replicate 1.** A-C) Plots of PCs 1-3 displaying the overlapping datasets. D) Plot of PC 1 and 2 of cells showing the variability of features associated with G1, S, G2M and NA phases. E) UMAP of S. cerevisiae labeled by cell cycle stages. Cells that could not be assigned to a cell cycle phase are marked as NA in D, E. F) Feature plots of histone genes enriched in S phase cells. G) UMAP plots of cells after cell-cycle regression, labeled by clusters obtained by unsupervised clustering. H) Stress response genes *DDR2*, *NCE103*, *HSP12*, and *TMA10* forming a separate cluster (cluster 2) after cell cycle regression. I) Feature plots of *AIM44*, *PIR1*, *PST1*, and *EGT2* for cell wall and budding marker expression that cluster separately (cluster 5) after cell cycle regression.

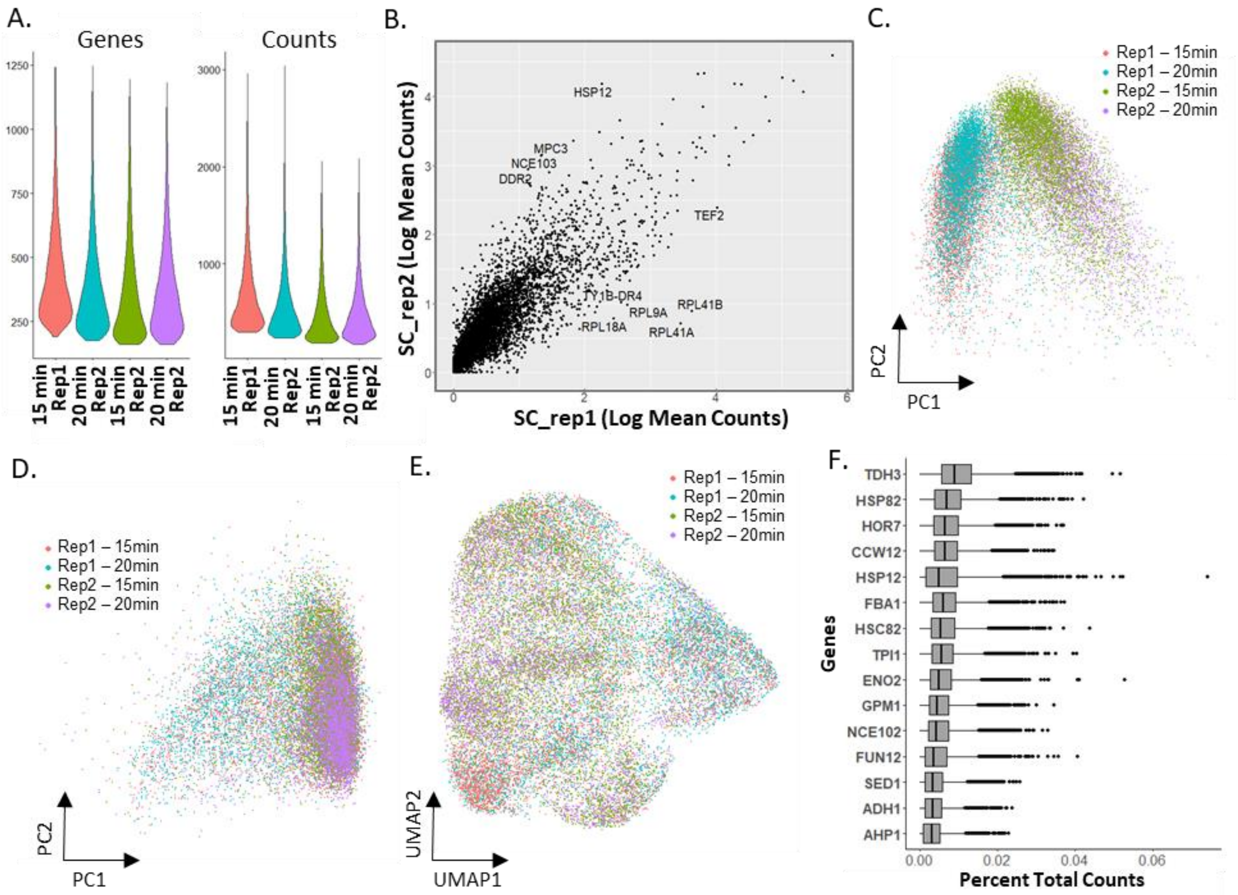


**Figure S2: Comparison of replicates for different *S. cerevisiae* lysis incubation time experiments.** A) The number of genes and UMI detected in 4 datasets. B) Plot of the average expression of genes in 22,491 *S. cerevisiae* cells from the control replicates. C-D) Plots of PC 1 and 2 displaying the four datasets without (C) and with (D) batch correction. E) UMAP plot of the datasets after batch correction. F) Top 15 genes expressed in the second replicate dataset.

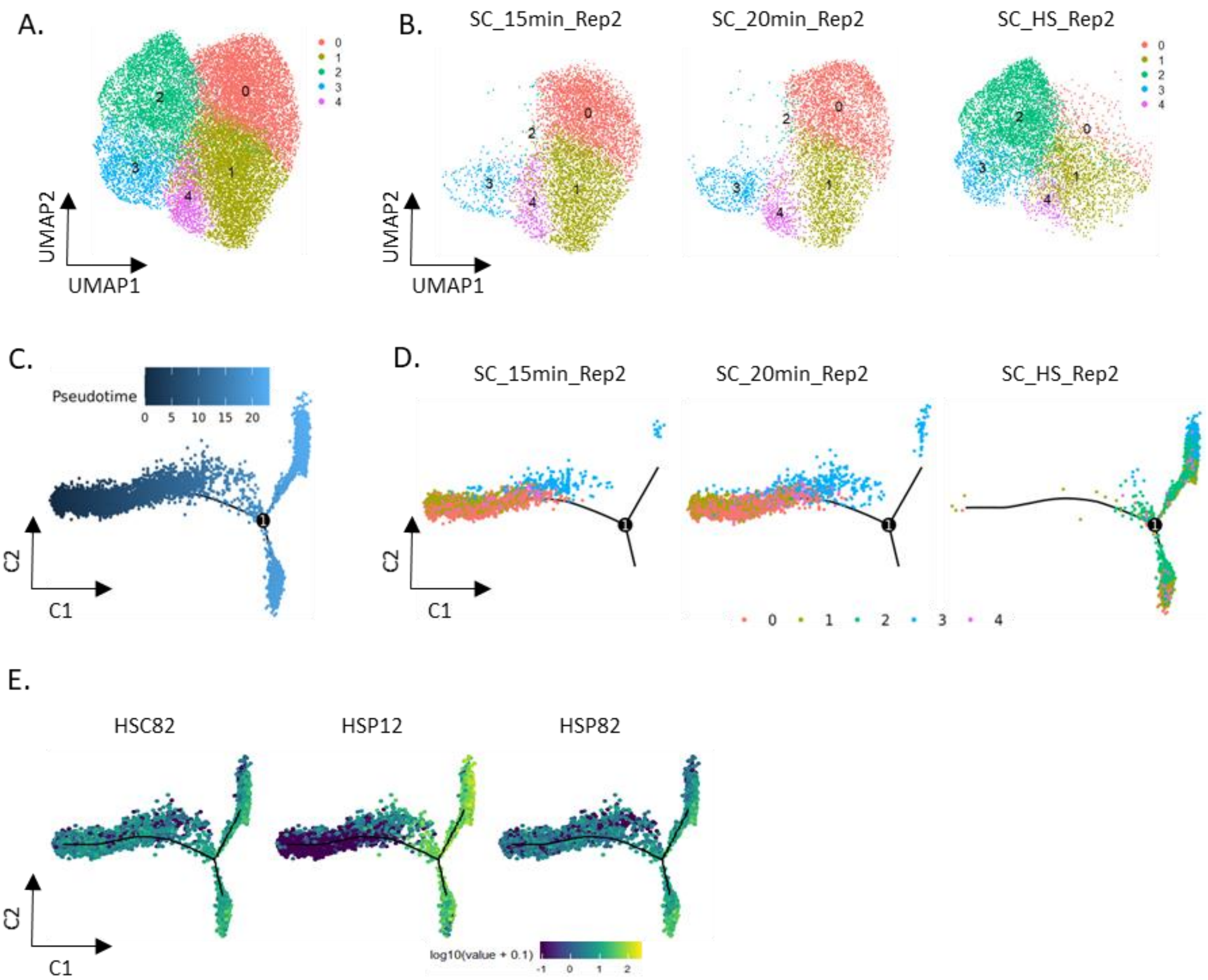


**Figure S3: Analysis of 26,019 *S. cerevisiae* cells in replicate 2 that combine control and heat-shock data, after regressing out cell-cycle effect.** A) UMAP plot of unsupervised clustering on the combined control (15 & 12 min lysis) and heat shocked *S. cerevisiae*, replicate 2 dataset after the cell cycle effects are regressed out. Colors represent cell clusters also shown in 2H. B) UMAP plots displaying the membership of cells from the controls (15 min-left; 20 min- middle) and heat-shocked (right) *S. cerevisiae* cells, replicate 2 shown in A. C) Inferred pseudo-time trajectory of the combined dataset. Color bar indicates pseudo-time. D) Inferred trajectories split by controls (15 min-left; 20 min- middle) and heat-shocked (right) datasets shown in Figure 2H. Colors represent cell clusters shown in Figures 2H, and S3A. E) Expression levels of heat shock genes *HSC82*, *HSP12* and *HSP82* along the pseudo-time trajectory. Color bar indicates scaled expression level.

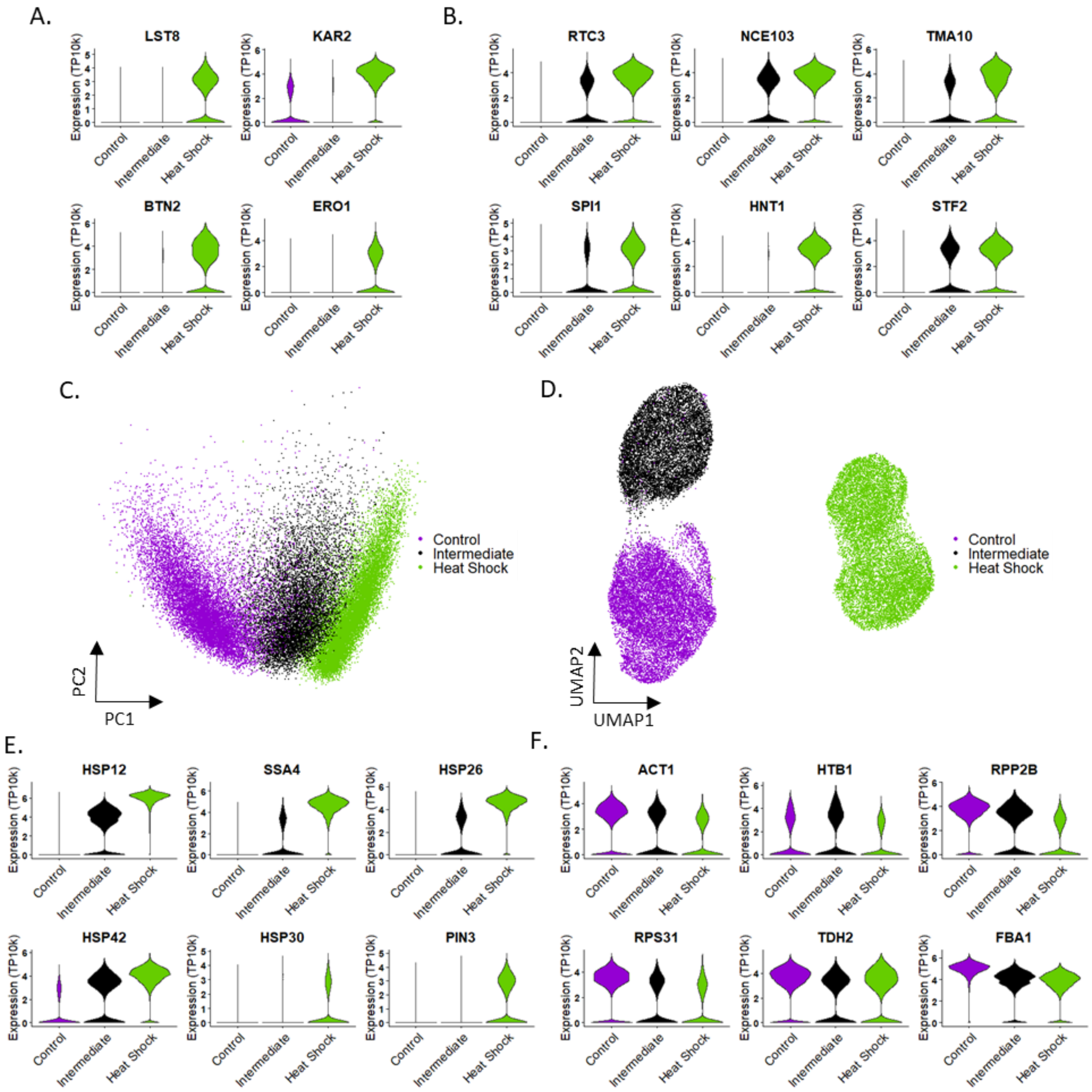


**Figure S4: Analysis of 35,109 *S. cerevisiae* cells including 14,007 cells after heat shock and 21,102 cells as control.** A) Four genes that indicate DNA replication stress show significant upregulation during heat shock when compared to the replicate 1 control. B) Several genes involved in protein transport through the ER and Golgi show elevated expression during heat shock. C) UMAP analysis of two heat-shock replicates and two control replicates without batch correction. D) UMAP analysis of two heat-shock replicates and only control replicate 2. E) Violin plots displaying heat-shock genes that are elevated in the second control replicate with yet higher expression in the heat-shock experiments. F) Violin plots displaying housekeeping genes (actin, histones, ribosomes, and glycolysis genes) that appear in both heat shock and control replicate 2 datasets.

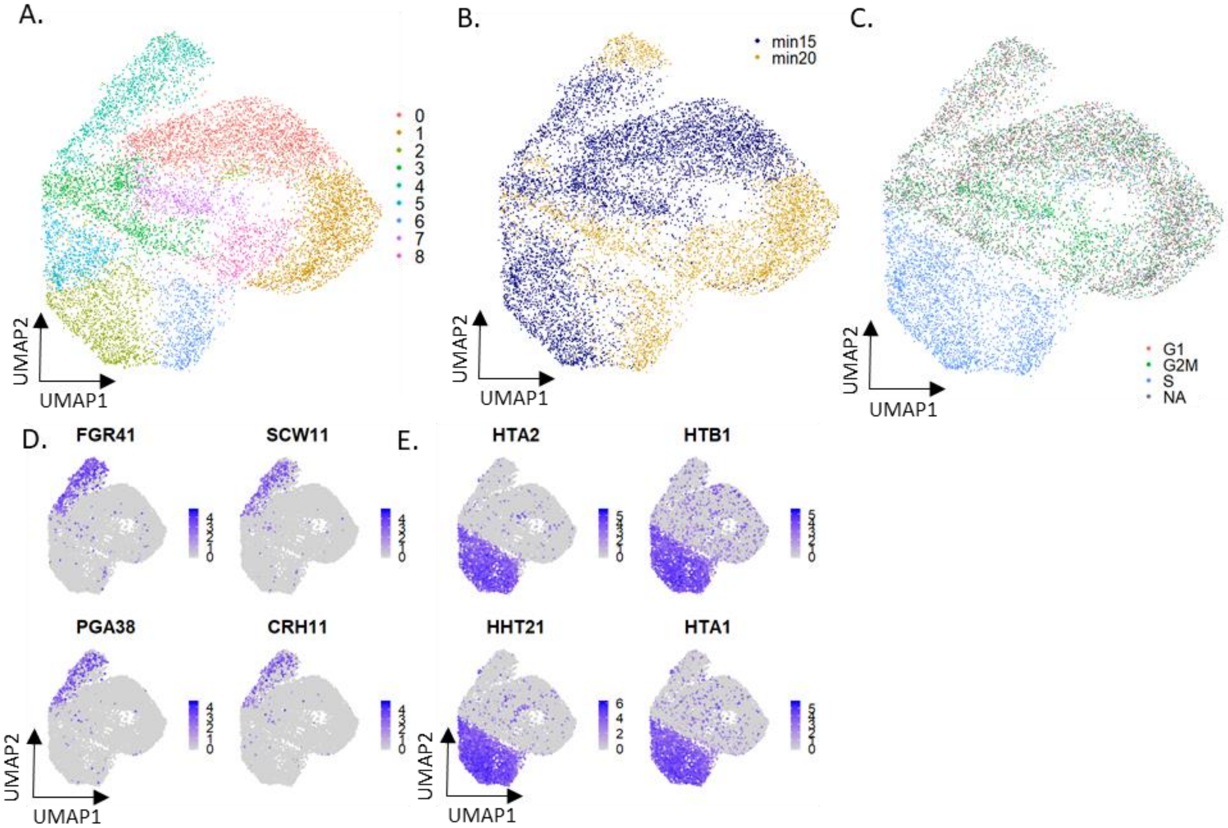


**Figure S5: mDrop-seq of 10,315 *C. albicans* cells from different lysis incubation times.** A) UMAP displaying the clustering analysis of 10,315 *C. albicans* cells detected. B) UMAP displaying how the different lysis time points cluster together. C) UMAP displaying cell cycle phase assignment across all cells. D) Feature plots displaying 4 GPI-anchored cell wall proteins that represent markers for cluster 4. E) Feature plots displaying 4 histone tail genes that represent markers for clusters 2, 5, and 6.

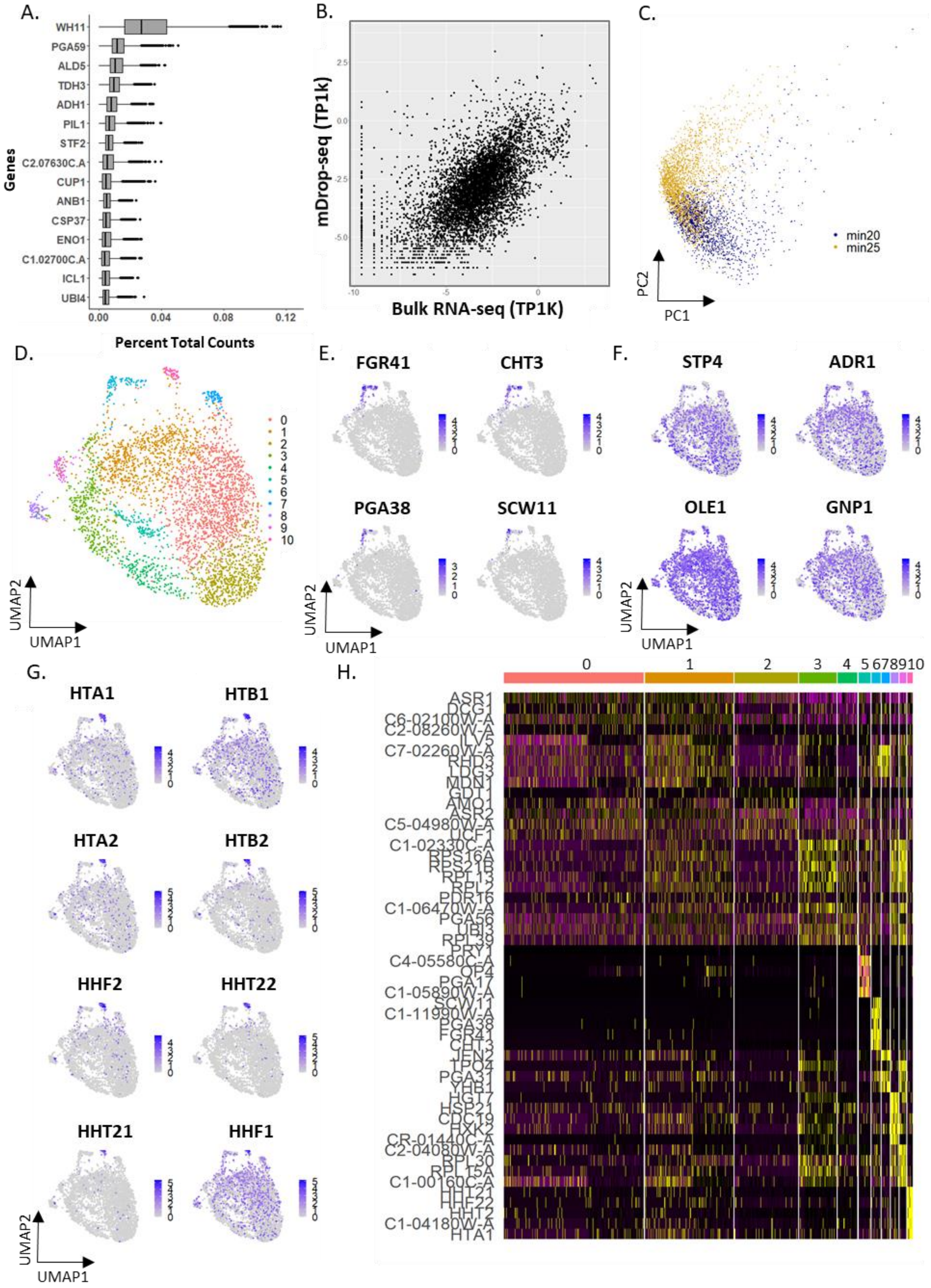

A.
WH11
PGA59
ALD5
TDH3
ADH1
PIL1
STF2
C2.07630C.A
CUP1
ANB1
CSP37
ENO1
C1.02700C.A
ICL1
UBI4
Genes
0.00
0.04
0.08
0.12
Percent Total Counts
B.
mDrop-seq (TP1k)
Bulk RNA-seq (TP1K)
C.
min20
min25
PC2
PC1
D.
0
1
2
3
4
5
6
7
8
9
10
UMAP2
UMAP1
E.
FGR41
CHT3
PGA38
SCW11
UMAP2
UMAP1
F.
STP4
ADR1
OLE1
GNP1
UMAP2
UMAP1
G.
HTA1
HTB1
HTA2
HTB2
HHF2
HHT22
HHT21
HHF1
UMAP2
UMAP1
H.
0
1
2
3
4
5
6
7
8
9
10
ASR1
C7-02260W-A
C1-02330C-A
C1-06470W-A
C4-05580C-A
C1-05890W-A
C1-11990W-A
PGA31
CR-01440C-A
C2-04080W-A
C1-00160C-A
C1-04180W-A
HTA1

**Figure S6: mDrop-seq of 4006 *C. albicans* cells in replicate experiments.** A) Boxplot of the 15 highest expression genes by percentage of total counts in the *C. albicans* data. B) A plot of average gene expression between CA_20min pseudo-bulk data and a Bulk RNA-seq dataset. C) A plot of PCs 1 and 2 looking at the lysis times 20 and 25 minutes. D) UMAP plot displaying the clustering analysis of the different *C. albicans* cells detected after batch correction. D) Feature plots displaying 4 cell wall genes that represent markers for cluster 6. F) Feature plots of transcription factors, fatty acid biosynthesis, and hyphal formation genes involved with *C. albicans* virulence. G) Feature plots displaying eight histone tail genes that represent markers for cluster 10. H) Heatmap displaying expressions of the top marker genes for each cluster shown in D.

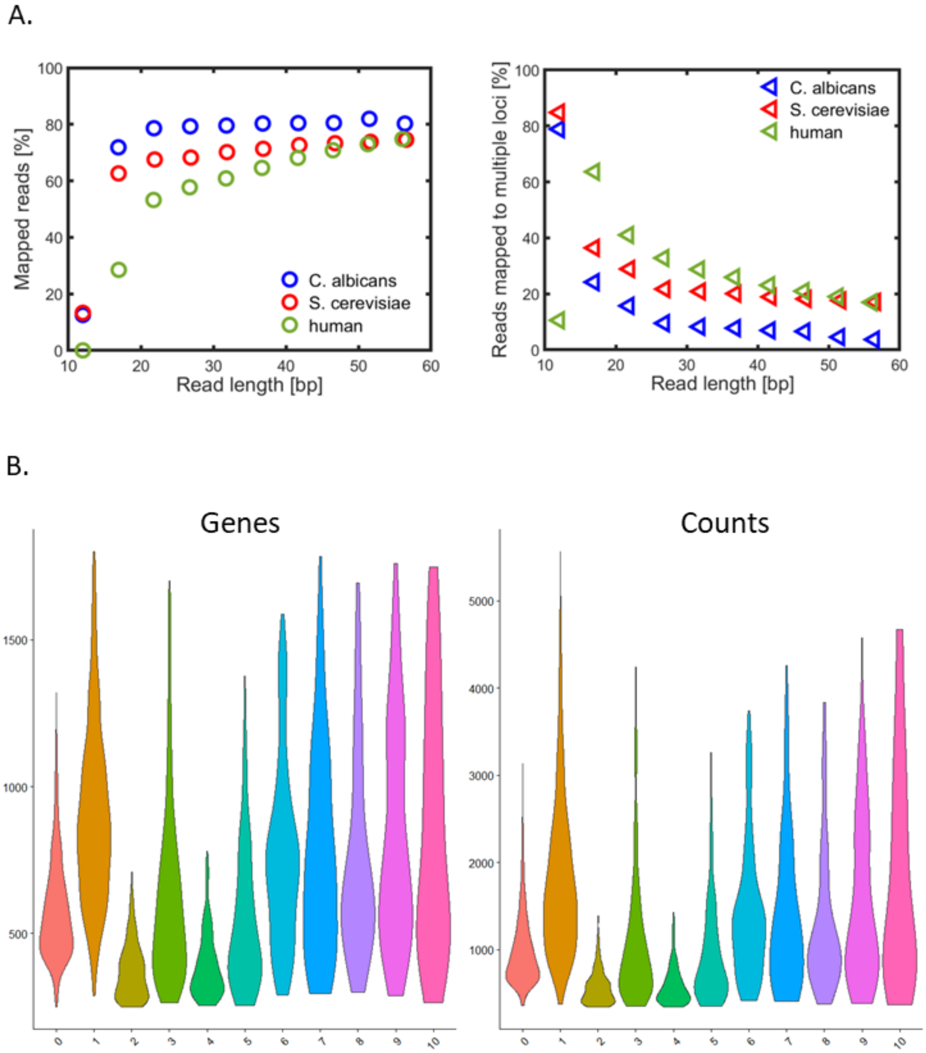


**Figure S7: Comparison of read lengths and mapping rates in *S. cerevisiae*, *C. albicans* and human scRNA-seq data.** A) Percent uniquely mapped reads (left) and reads mapped to multiple loci (right) as function of read length, for *C. albicans*, *S. cerevisiae* and human genomes. B) *C. albicans* expression levels across clusters in Figure S6D. Clusters 6-10, the separate cluster containing histone genes, glycolysis genes, and cell wall genes, show slightly higher rate of genes or UMI detected compared to other clusters.

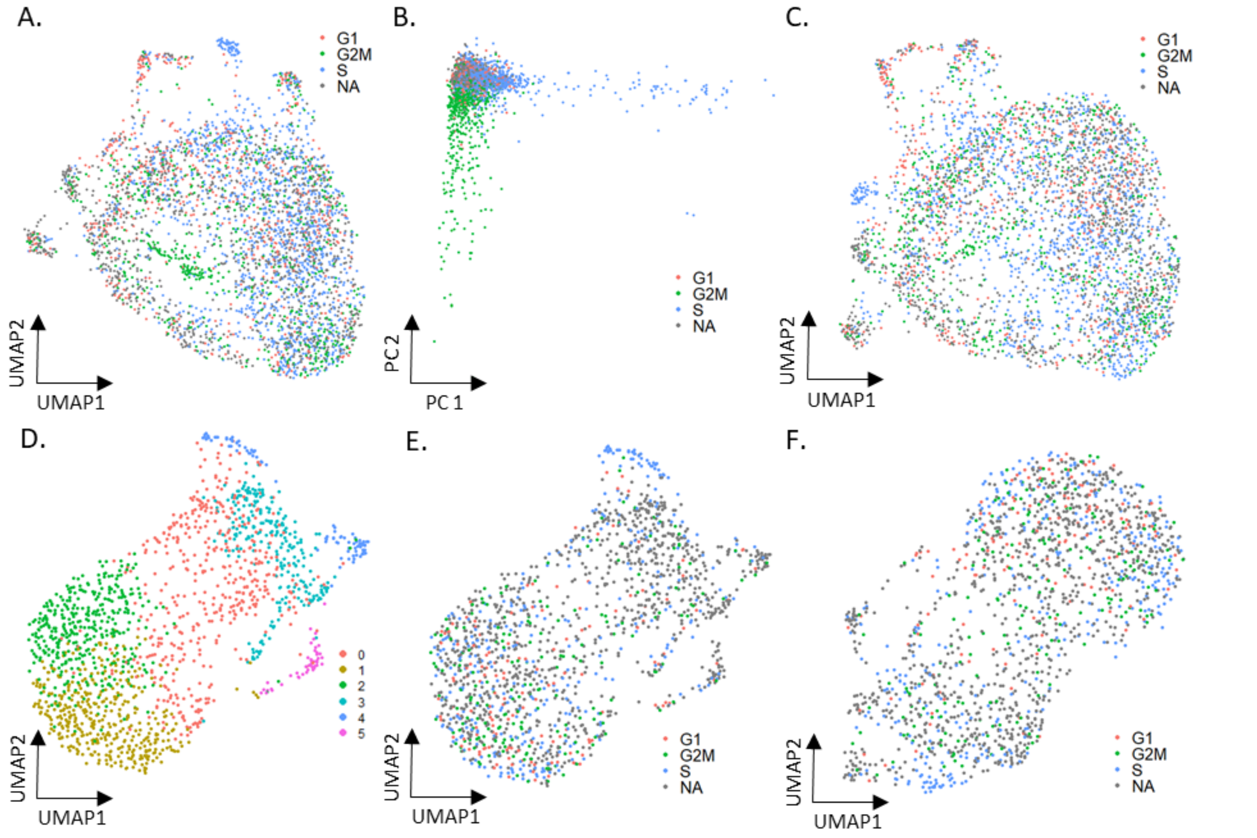


**Figure S8: Analysis of cell cycle genes in *C. albicans* replicates**. A) UMAP plot from Figure S6D with cells marked for the G1, G2M and S phases of the cell cycle. B) Plot of PC 1 and 2 showing the variation of cell cycle genes in the integrated dataset. C) UMAP plot showing the integrated dataset after regressing out the variations from cell cycle. D) UMAP plot of CA_rep1_20min dataset only, with cells colored by clusters. E) UMAP plot of CA_rep1_20min dataset with cells colored by their cell cycle scoring. F) UMAP plot of the CA_rep1_20min dataset with variations from cell cycle regressed out.

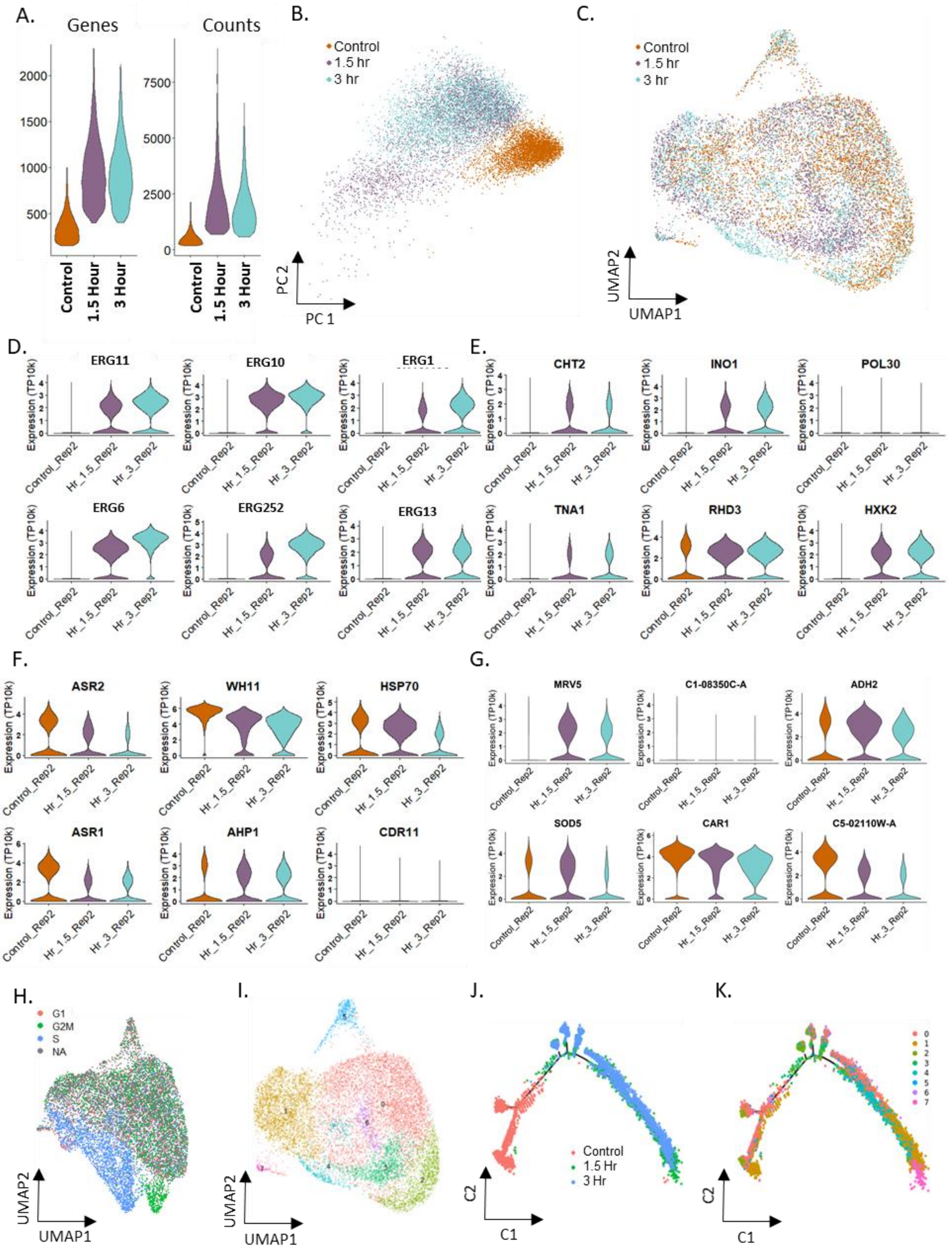
A.
Genes
Counts
Control
1.5 Hour
3 Hour
B.
Control
1.5 hr
3 hr
PC 2
PC 1
C.
Control
1.5 hr
3 hr
UMAP2
UMAP1
D.
ERG11
ERG10
ERG1
ERG6
ERG252
ERG13
E.
CHT2
INO1
POL30
TNA1
RHD3
HXK2
F.
ASR2
WH11
HSP70
ASR1
AHP1
CDR11
G.
MRV5
C1-08350C-A
ADH2
SOD5
CAR1
C5-02110W-A
Expression (TP10k)
Control_Rep2
Hr_1.5_Rep2
Hr_3_Rep2
H.
G1
G2M
S
NA
UMAP2
UMAP1
I.
UMAP2
UMAP1
J.
Control
1.5 Hr
3 Hr
C2
C1
K.
C2
C1

**Figure S9: Fluconazole treatment of 10,942 *C. albicans* cells, replicate 2 profiled using mDrop-seq.** A) Violin plot displaying the number of genes and UMI for each heat shock replicate. B) Plot of PCs 1 and 2 displaying the control and fluconazole datasets. C) UMAP displaying the clustering patterns of the integrated control and fluconazole libraries. D) Violin plots displaying the expression differences in control and fluconazole datasets for several ergosterol biosynthesis genes. E) UMAP displaying cell cycle assignment in combined dataset containing control and fluconazole treated cells. F) UMAP of cell clusters obtained from unsupervised clustering of combined dataset after cell cycle effects are regressed out. G-I) Violin plots of genes that show differences in expression between control and fluconazole treated datasets. G) Genes where RNA levels decrease progressively between control and fluconazole treated cells, 1.5 and 3 hr treatment. H) Genes where RNA levels increase progressively with increased fluconazole exposure. I) Genes from the intermediate treatment timepoint, 1.5 hr that show higher expression compared to control and 3 hr fluconazole treated cells. J, K) Pseudo-time trajectory of the combined dataset inferred using Monocle. Colors indicate J) experimental time points, and K) cell-type clusters shown in I.

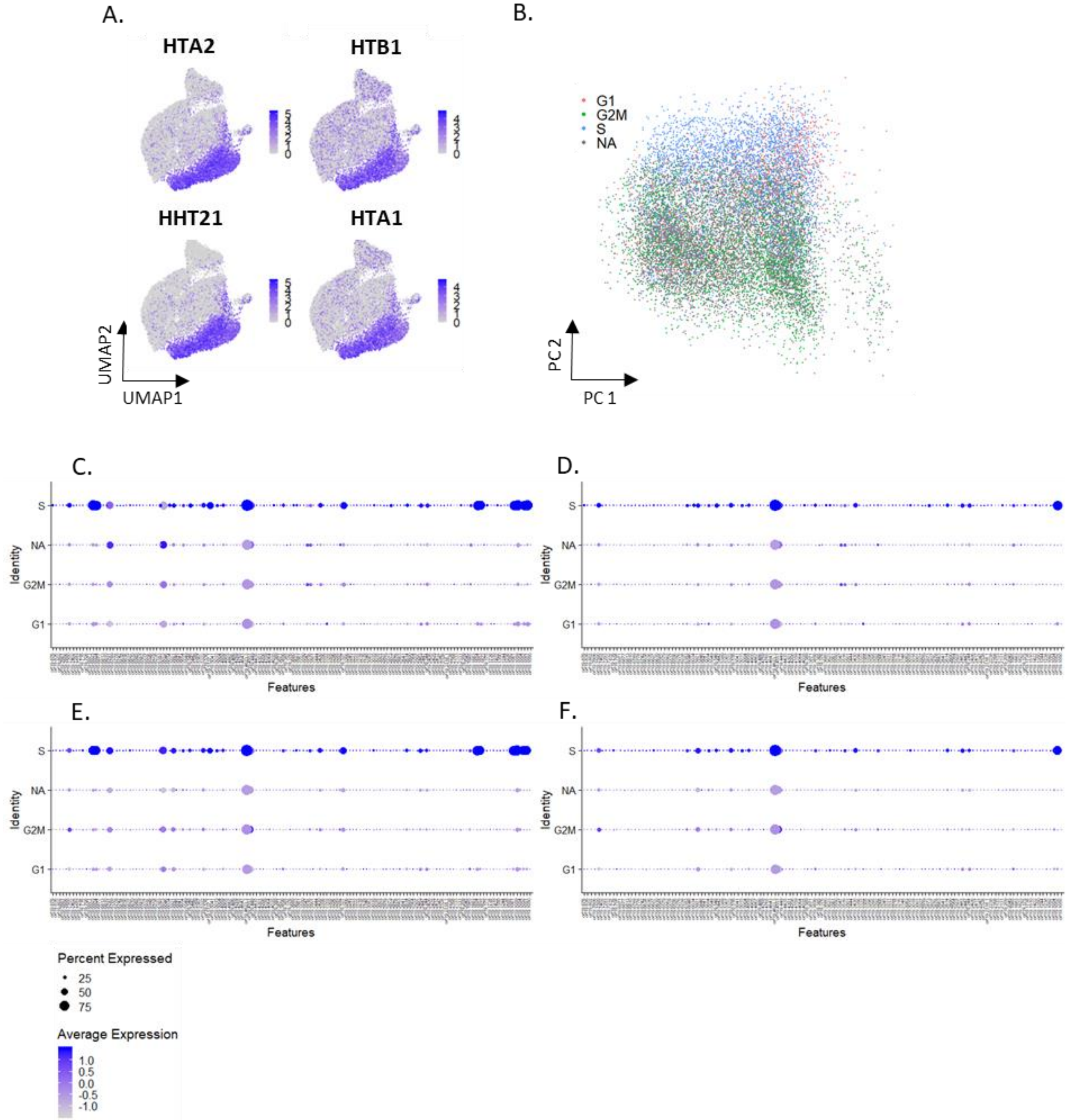


**Figure S10: Cell cycle analysis of 15,503 control and Fluconazole treated *C. albicans* cells, replicate 1.** A) Feature plots displaying 4 histone tail genes across control and fluconazole exposed cells. B) Plot of PCs 1 and 2 featuring the variability of features associated with S and G2M gene markers. C-F) Dot plots of S phase markers C) with histone genes on replicate 1; D) without histone genes on replicate 1; E) with histone genes on replicate 2; F) without histone genes on replicate 2.

258

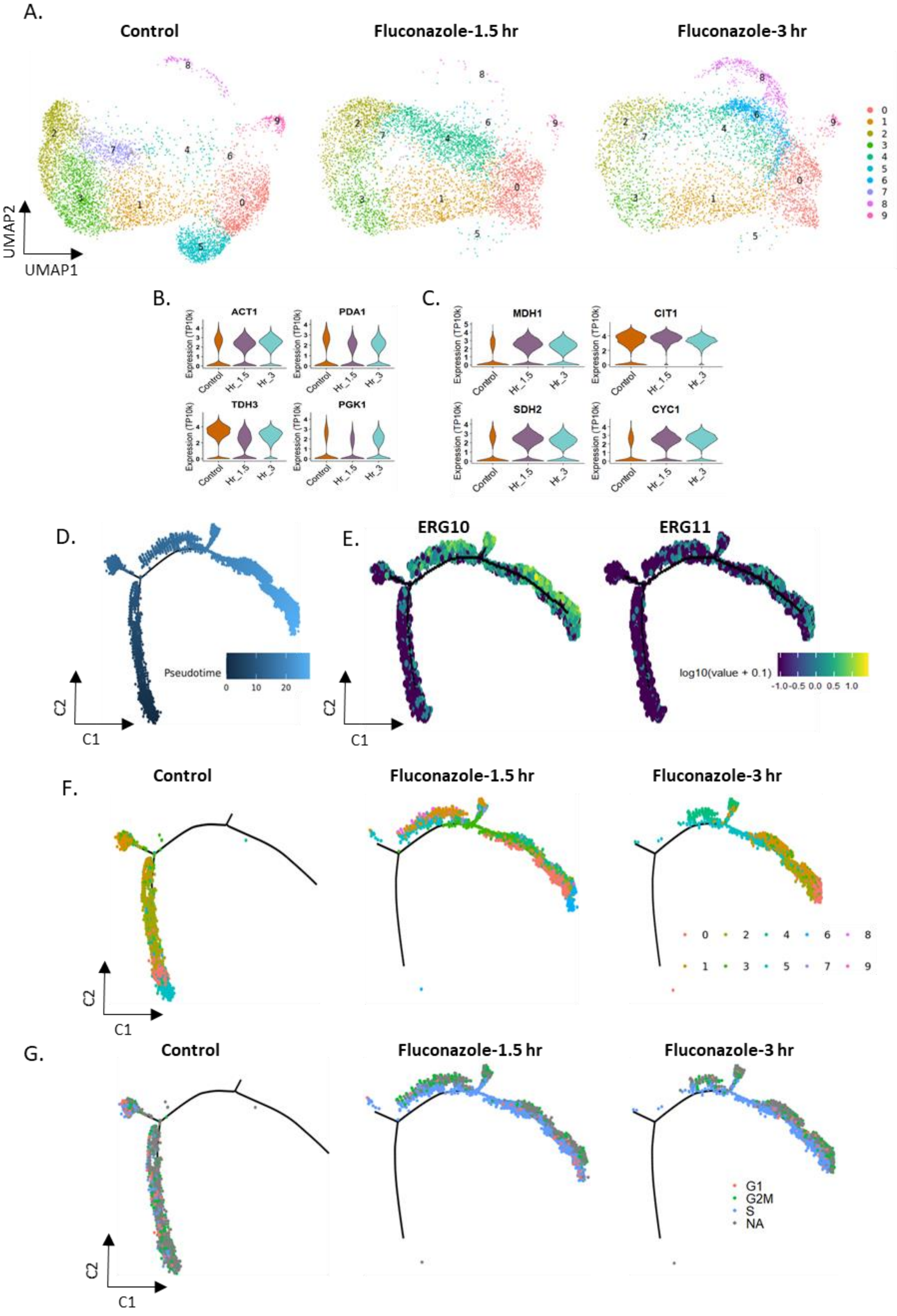
A.
Control
Fluconazole-1.5 hr
Fluconazole-3 hr
UMAP2
UMAP1
B.
ACT1
PDA1
TDH3
PGK1
C.
MDH1
CIT1
SDH2
CYC1
Expression (TP10k)
Control
Hr_1.5
Hr_3
D.
E.
ERG10
ERG11
Pseudotime
log10(value + 0.1)
C2
C1
F.
Control
Fluconazole-1.5 hr
Fluconazole-3 hr
G.
Control
Fluconazole-1.5 hr
Fluconazole-3 hr
G1
G2M
S
NA

**Figure S11: Fluconazole treatment of 15,503 *C. albicans*, replicate 1.** A) UMAP plots displaying the membership of cells from the control (left) and fluconazole (1.5 hr- middle; 3 hr- right) samples in the combined dataset shown in Figure 4I. B) Expression levels of housekeeping genes *ACT1*, *PDA1*, *TDH3* and *PGK1* in replicate 1. C) Expression levels of *MDH1*, *CIT1*, *SDH2*, *CYC1* genes that are differentially expressed in cluster 5 in A and Figure 4I. D) Inferred pseudo-time trajectory of the combined dataset. Color bar indicates pseudo-time. E) Expression levels of ergosterol synthesis genes *ERG10* and *ERG11* increase with pseudo-time. F-G) Inferred trajectories split by control (left) and fluconazole (1.5 hr- middle; 3 hr- right) datasets shown in Figure 4K. Colors represent F) cell clusters shown in Figure 4I, G) Cell cycle phases, G1, S, G2M and not assigned (NA).

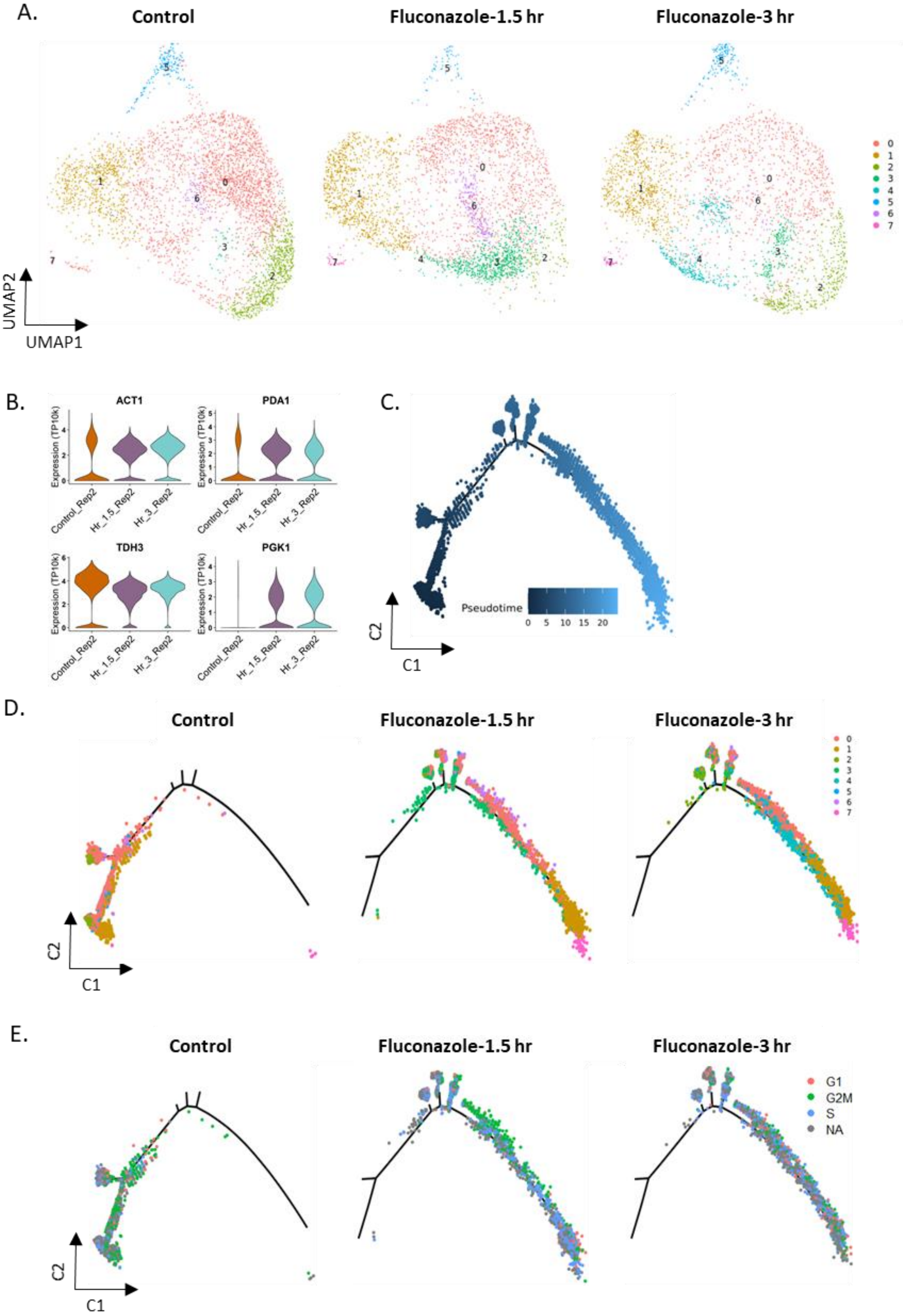
A.
Control
Fluconazole-1.5 hr
Fluconazole-3 hr
UMAP2
UMAP1
B.
ACT1
PDA1
TDH3
PGK1
Expression (TP10k)
Control_Rep2
Hr_1.5_Rep2
Hr_3_Rep2
C.
Pseudotime
0 5 10 15 20
C2
C1
D.
Control
Fluconazole-1.5 hr
Fluconazole-3 hr
C2
C1
E.
Control
Fluconazole-1.5 hr
Fluconazole-3 hr
G1
G2M
S
NA
C2
C1

**Figure S12: Fluconazole treatment of 10,942 *C. albicans*, replicate 2.** A) UMAPs display the membership of cells from the control (left) and fluconazole (1.5 hr- middle; 3 hr- right) samples in the combined datasets shown in Figure S9F. B) Expression levels of common housekeeping genes *ACT1*, *PDA1*, *TDH3*, *PGK1* in replicate 2. C) Inferred trajectory, with color bar indicating pseudo-time. D-E) Trajectories split by control (left) and fluconazole (1.5 hr- middle; 3 hr- right) datasets in Figure S9H. Colors represent D) cell clusters shown in Figure S9F, E) Cell cycle phases G1, S, G2M and not assigned (NA).

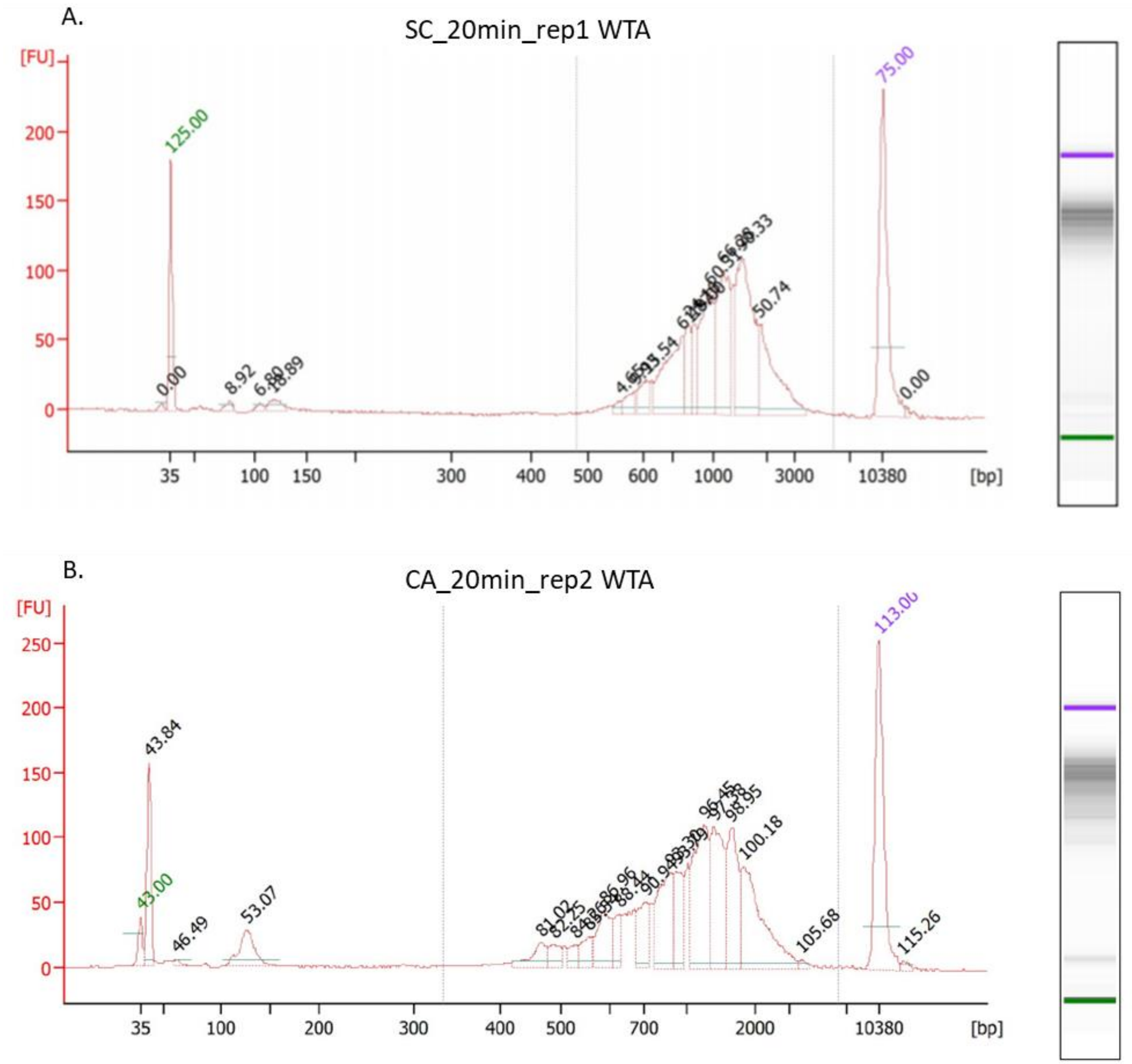


**Figure S13: Distribution of barcoded cDNA obtained from mDrop-seq experiments after Whole Transcriptome Amplification (WTA).** A) *S cerevisiae* (SC_20min_Rep1) and B) *C. albicans* (CA_20min_Rep2), shown in Agilent Bioanalyzer traces.

**Table S1:** Summary of *S. cerevisiae and C. albicans* datasets showing average read counts, genes, and UMI detected per cell for mDrop-seq experiments. The average reads and counts for the species mixing experiments are reported after filtering out misaligned genes.

| Library | Raw Counts | Cells Detected | Average Counts per cell | Average Genes per cell | Median Genes per cell | Average UMI per cell | Median UMI per cell |
|---|---|---|---|---|---|---|---|
| SC_15min_Rep1 | 227,973,901 | 6,620 | 34,437 | 474 | 417 | 781 | 639 |
| SC_20min_Rep1 | 137,063,719 | 5,392 | 25,419 | 402 | 347 | 626 | 502 |
| SC_15min_Rep2 | 108,347,980 | 4,350 | 24,907 | 380 | 334 | 503 | 424 |
| SC_20min_Rep2 | 58,362,030 | 4,740 | 12,312 | 404 | 365 | 529 | 460 |
| SC_Heat Shock_Rep1 | 65,393,958 | 7,483 | 8,739 | 434 | 377 | 721 | 590 |
| SC_Heat Shock_Rep2 | 54,662,686 | 6,524 | 8,740 | 587 | 515 | 1,053 | 866 |
| Species Mix- SC - 7% | 88,687,370 | 1,495 | 21,288 | 544 | 487 | 1,108 | 911 |
| Species Mix- CA - 7% | 88,687,370 | 2,671 | 21,288 | 193 | 137 | 422 | 348 |
| Species Mix - SC - 15% | 88,953,431 | 4,901 | 10,962 | 578 | 529 | 1,173 | 981 |
| Species Mix - CA - 15% | 88,953,431 | 3,214 | 10,961 | 797 | 717 | 1,565 | 1,276 |
| CA_15min_Rep1 | 30,400,506 | 1,215 | 25,021 | 451 | 376 | 685 | 504 |
| CA_20min_Rep1 | 93,129,841 | 1,693 | 55,008 | 837 | 706 | 1,660 | 1,223 |
| CA_25min_Rep1 | 128,016,172 | 2,364 | 54,152 | 573 | 494 | 1,029 | 783 |
| CA_15min_Rep2 | 96,778,937 | 6,452 | 7,132 | 508 | 549 | 836 | 610 |
| CA_20min_Rep2 | 129,649,186 | 3,862 | 15,021 | 611 | 561 | 1,132 | 976 |
| CA_Fluconazole-ctrl_Rep1 | 154,646,838 | 5,952 | 25,982 | 563 | 510 | 944 | 784 |
| CA_Fluconazole-1.5hr_Rep1 | 122,638,009 | 4,924 | 24,906 | 893 | 816 | 1,948 | 1586 |
| CA_Fluconazole-3hr_Rep1 | 111,642,039 | 4,627 | 24,128 | 953 | 922 | 2,130 | 1,882 |
| CA_Fluconazole-ctrl_Rep2 | 95,442,703 | 4,227 | 10,918 | 351 | 324 | 527 | 456 |
| CA_Fluconazole-1.5hr_Rep2 | 115,687,150 | 3,687 | 12,978 | 967 | 918 | 2,152 | 1,863 |
| CA_Fluconazole-3hr_Rep2 | 96,686,038 | 3,028 | 11,989 | 935 | 896 | 1,909 | 1,689 |

**Table S2:** The number and percentage of cells in different cell cycle phases across the different experimental replicates.

| Dataset | G1 Cells | G1 (%) | S Cells | S (%) | G2M Cells | G2M (%) | NA Cells | NA (%) |
|---|---|---|---|---|---|---|---|---|
| SC_Rep1 | 1,194 | 15.8 | 3,435 | 45.4 | 2,938 | 38.8 | 1,523 | 16.7 |
| SC_Rep2 | 2,453 | 27.5 | 3,615 | 40.5 | 2,862 | 32 | 3,082 | 25.7 |
| SC_Heat Shock _Rep1 | 1,016 | 18.4 | 3,770 | 68.2 | 738 | 13.4 | 1,369 | 19.9 |
| SC_Heat Shock _Rep2 | 831 | 15.3 | 3,636 | 66.8 | 979 | 18 | 1,078 | 16.5 |
| CA_Rep1 | 268 | 14.6 | 862 | 47.1 | 702 | 38.3 | 2,174 | 54.3 |
| CA_Rep2 | 671 | 11.4 | 3,282 | 55.8 | 1,932 | 32.8 | 4,429 | 42.9 |
| CA_Fluconazole-ctrl_Rep1 | 585 | 25.3 | 702 | 30.3 | 1,029 | 44.4 | 3,636 | 61.1 |
| CA_Fluconazole-1.5hr_Rep1 | 370 | 11.7 | 2,257 | 71.7 | 522 | 16.6 | 1,775 | 36 |
| CA_Fluconazole-3hr_Rep1 | 603 | 22.2 | 1,325 | 48.9 | 782 | 28.9 | 1,917 | 41.4 |
| CA_Fluconazole-ctrl_Rep2 | 351 | 17.9 | 756 | 38.6 | 854 | 43.5 | 2,266 | 53.6 |
| CA_Fluconazole-1.5hr_Rep2 | 233 | 10.2 | 1,344 | 58.8 | 708 | 31 | 1,402 | 38 |
| CA_Fluconazole-3hr_Rep2 | 273 | 16.6 | 927 | 56.3 | 447 | 27.1 | 1,381 | 45.6 |
